# Chronic inorganic nitrate supplementation does not improve metabolic health and worsens disease progression in mice with diet-induced obesity

**DOI:** 10.1101/2024.07.04.602070

**Authors:** Alice P. Sowton, Lorenz M.W. Holzner, Fynn N. Krause, Ruby Baxter, Gabriele Mocciaro, Dominika K. Krzyzanska, Magdalena Minnion, Katie A. O’Brien, Matthew C. Harrop, Paula M. Darwin, Benjamin D. Thackray, Michele Vacca, Martin Feelisch, Julian L. Griffin, Andrew J. Murray

## Abstract

Inorganic nitrate (NO_3_^-^) has been proposed to be of therapeutic use as a dietary supplement in obesity and related conditions including the Metabolic Syndrome (MetS), type-II diabetes and metabolic dysfunction associated steatotic liver disease (MASLD). Administration of NO_3_^-^ to endothelial nitric oxide synthase-deficient mice reversed aspects of MetS, however the impact of NO_3_^-^ supplementation in diet-induced obesity is not well understood. Here we investigated the whole-body metabolic phenotype and cardiac and hepatic metabolism in mice fed a high-fat high-sucrose (HFHS) diet for up to 12-months of age, supplemented with 1 mM NaNO_3_ (or NaCl) in their drinking water. HFHS-feeding was associated with a progressive obesogenic and diabetogenic phenotype, which was not ameliorated by NO_3_^-^. Furthermore, HFHS-fed mice supplemented with NO_3_^-^ showed elevated levels of cardiac fibrosis, and accelerated progression of MASLD including development of hepatocellular carcinoma in comparison with NaCl-supplemented mice. NO_3_^-^ did not enhance mitochondrial β-oxidation capacity in any tissue assayed and did not suppress hepatic lipid accumulation, suggesting it does not prevent lipotoxicity. We conclude that NO_3_^-^ is ineffective in preventing the metabolic consequences of an obesogenic diet and may instead be detrimental to metabolic health against the background of HFHS-feeding. This is the first report of an unfavorable effect of long-term nitrate supplementation in the context of the metabolic challenges of overfeeding, warranting urgent further investigation into the mechanism of this interaction.

**New & Noteworthy:** Inorganic nitrate has been suggested to be of therapeutic benefit in obesity-related conditions as it increases nitric oxide bioavailability, enhances mitochondrial β-oxidation and reverses Metabolic Syndrome in *eNOS^-/-^* mice. However, we here show that over 12 months, nitrate was ineffective in preventing metabolic consequences in high-fat high-sucrose fed mice, and worsened aspects of metabolic health, impairing cholesterol handling, increasing cardiac fibrosis, and exacerbating steatotic liver disease progression, with acceleration to hepatocellular carcinoma.

## Introduction

The obesity epidemic remains a global health concern, and its prevalence continues to increase worldwide (1). Obesity is associated with high morbidity and mortality (2), largely due to the elevated risk of comorbidities such as type-II diabetes mellitus (T2DM), cardiovascular disease (CVD) and metabolic dysfunction associated steatotic liver disease (MASLD), a condition previously known as non-alcoholic fatty liver disease (NAFLD) (3–7).

Obesity and diabetes are associated with hypertension (8), which is in turn associated with endothelial dysfunction and reduced nitric oxide (NO) bioavailability. Mice lacking endothelial NO synthase (*eNOS^-/-^*mice) are hypertensive (9), but also develop symptoms of the Metabolic Syndrome (MetS) including glucose intolerance and dyslipidemia, alongside hyperleptinemia (10, 11) and defective mitochondrial β-oxidation and biogenesis (12, 13). In humans, eNOS polymorphisms have been associated with T2DM and MetS (14, 15), whilst patients with T2DM generate less NO from L-arginine than healthy controls (16). NO production and metabolism, particularly decreased NO bioavailability, is therefore considered central to the etiology of MetS/T2DM.

Canonical formation of NO occurs via oxidation of a guanidino nitrogen of L-arginine by one of 3 isoforms of NOS. However, it is now recognized that NO can also be produced via reduction of endogenously produced or dietary inorganic nitrate (NO_3_^-^; [17]). In the circulation, NO generated by the vascular endothelium is oxidized to nitrite (NO_2_^-^) and nitrate. The latter is taken up from the blood and secreted into saliva where it is reduced to nitrite by commensal bacterial flora of the mouth (18). Upon swallowing of saliva, NO_2_^-^ is rapidly protonated in the stomach forming nitrous acid (HNO_2_), which spontaneously decomposes to form NO (19, 20). Remaining nitrate or nitrite can be absorbed in the intestine, where nitrite can be reduced to NO by enzymes including xanthine oxidoreductase, deoxyhemoglobin, and myoglobin (21–23). Circulating NO_3_^-^ is eventually excreted by the kidneys, although ∼25% is actively taken up by the salivary glands allowing for it to be concentrated in saliva and recirculated (17, 20). Thus, inorganic nitrate, once considered an end product of NO metabolism (24) and, along with nitrite, a potentially toxic residue of food preservation (25), is now recognized as a potentially important source of NO under hypoxic and acidotic conditions (20, 26) and a route to modulate NO bioavailability via dietary manipulation (20). Both nitrite and nitrate were used medicinally long before any of these mechanisms had been discovered (27).

Dietary supplementation with inorganic nitrate for 8-10 weeks reverses features of MetS in *eNOS^-/-^* mice, improving glucose handling, hypertension, and dyslipidemia (28). It is not established whether reduced NO bioavailability is a universal feature of MetS and/or T2DM, however nitrite improved glycaemia in *db/db* mice (one month treatment; [29]) and *ob^lep^* mice (one week treatment; [30]), whilst nitrate improved insulin sensitivity in high-fructose-fed rats over 10 weeks (31), high-fat diet low-dose streptozotocin T2DM rats over two months (32) and high-fat-high-fructose-fed mice over one month (33). At a tissue level, dietary nitrate increased mitochondrial β-oxidation in the heart and skeletal muscle of rats and mice (34–36), and was associated with increased voluntary wheel running in mice (37). Dietary inorganic nitrate enhanced white adipose tissue browning in rats (38), and increased mitochondrial oxygen consumption (39) and glucose oxidation (40) in white adipocytes. In liver, NO (generated by *eNOS*) inhibits activation of pro-inflammatory Kupffer cells, a response characteristic of MASLD pathogenesis (41). Although, long-term (17 months) nitrate supplementation did not result in adverse health effects in healthy mice and was associated with improved insulin sensitivity (42), the long-term implications of nitrate supplementation in obese animals remain unknown. This could be of particular importance in light of the effects of overfeeding with lipids and carbohydrates on mitochondrial function and reactive oxygen species (ROS) production against the complex interaction with tissue and whole-body redox regulation *vis-a-vis* aging-related alterations in metabolic regulation by liver and skeletal muscle.

We therefore sought to investigate whether dietary supplementation with a moderate dose of inorganic nitrate (similar to that achievable with a human diet rich in leafy vegetables) ameliorates the progression of metabolic comorbidities associated with diet-induced obesity in mice. We hypothesized that inorganic nitrate would delay the development of metabolic and mitochondrial dysregulation in high-fat high-sucrose-fed mice via enhanced NO bioavailability and modulation of tissue mitochondrial function.

## Materials and Methods

### Chemicals

Unless otherwise stated, all reagents were purchased from Sigma-Aldrich (Merck Life Science UK, Gillingham, UK).

### Ethical Approval

All studies were carried out in accordance with United Kingdom Home Office legislation under the Animals in Scientific Procedures (1986) Act and received prior approval from the University of Cambridge Animal Welfare and Ethical Review Board. All procedures were carried out by a personal license holder in accordance with these regulations.

### Study Design

Male C57Bl/6J mice (n = 95; RRID: MGI:3028467) were purchased from a commercial breeder (Charles River Laboratories, Margate, UK) at 3 weeks of age. Unless otherwise stated, all mice were group-housed in conventional cages under controlled environmental conditions (21°C, 54% humidity, 12 h photoperiod) and were allowed *ad libitum* access to food and water for the duration of the study.

In the first week of vivarium acclimatization, mice received a standard laboratory rodent chow diet (RM3(E), Special Diet Services, Essex, UK; 3.6 kcal.g^-1^, 39.7% carbohydrate, 22.4% crude protein, 4.3% crude fat, forthwith referred to as ‘chow’) and distilled water *ad libitum*. From 4 weeks of age, mice were randomized into four experimental groups (n=23-24/group); two groups continued to receive chow, whilst the others received a high-saturated fat, high-sucrose (HFHS) diet (TD.88137, Envigo Teklad Diets, Madison, WI, USA; 4.5 kcal.g^-1^, 48.5% carbohydrate, 17.3% crude protein, 21.2% crude fat). Within each diet group, one experimental group received drinking water supplemented with 1 mM sodium nitrate (NaNO_3_; TraceSELECT™, #71752, Fluka; Honeywell Specialty Chemicals, Seelze, Germany) whilst the other received water supplemented with 1 mM sodium chloride (NaCl; TraceSELECT™, #38979, Fluka) to control for sodium intake. Mice were maintained on their respective diets/treatments until they reached 4-, 8- or 12-months of age (n=7-8/group, **Fig. 1A**). Body mass, food and water intake were measured weekly.

**Figure 1.**
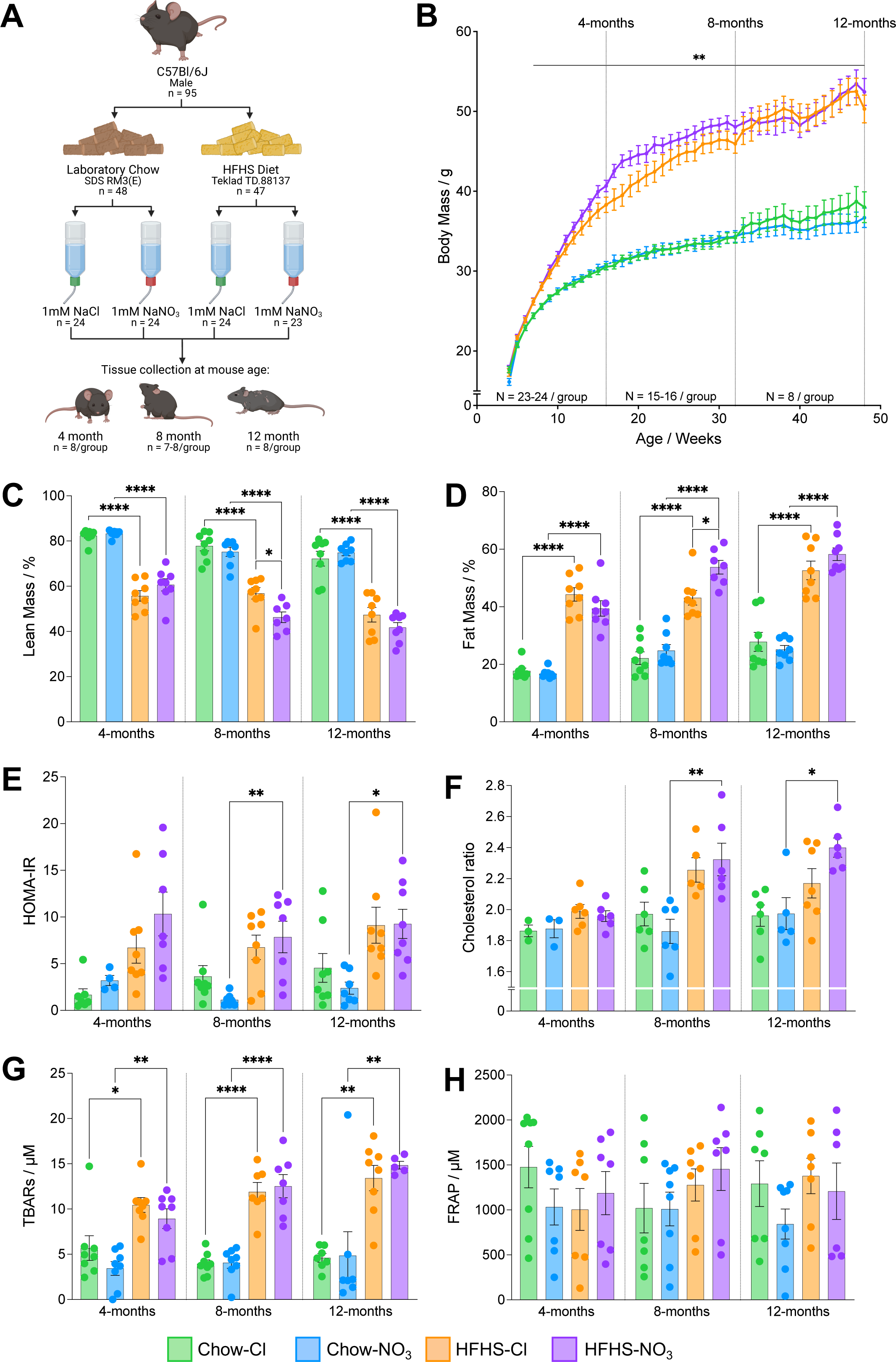
Mice fed a high-fat high-sucrose diet show progressive diet-induced obesity. (A) Experimental design. Male C57Bl/6J mice were randomly allocated to receive either standard laboratory chow (‘Chow’; Special Diet Services RM3 (E)) or a high-fat high-sucrose diet (‘HFHS’; Teklad TD.88137) from 4-weeks of age. Simultaneously, mice were randomized to receive either sodium chloride (NaCl; 1 mM) or sodium nitrate (NaNO_3_; 1 mM) *via* their drinking water. Mice continued to receive these dietary interventions *ad libitum* until they reached 4-, 8- or 12-months of age, where tissues were collected for subsequent metabolic analyses (n=7-8/group). Created with BioRender.com. (B) Mouse body mass over the course of the study. Individual mouse body mass was measured once per week. **p<0.01; HFHS compared with chow-fed groups supplemented with the same water treatment, for each week (two-way ANOVA with Benjamini-Hochberg FDR correction and Tukeys’ *post hoc* HSD test). (C) Percentage (%) of the body carcass (following removal of the heart and liver) that represents lean mass determined via dual-energy x-ray absorptiometry. N=7-8 / group. (D) Percentage (%) of the body carcass (following removal of the heart and liver) that represents fat mass determined via dual-energy x-ray absorptiometry. N=7-8 / group. (E) Homeostatic model of insulin resistance calculated according to (48) using measurements of fasted blood glucose and insulin. N=4-8 / group. (F) Cholesterol ratio (total cholesterol / HDL cholesterol), used clinically as a predictor of cardiovascular disease (62), calculated from clinical biochemistry measurements. N=3-7 / group. (G) Concentration of thiobarbituric acid reactive substances (TBARS) in plasma indication levels of lipid peroxidation. N=5-8/group. (H) Ferric reducing ability of plasma (FRAP), as an indication of the antioxidant capacity of plasma. (N 6-8 / group). For (C)-(H) data represent mean ± SEM, *p<0.05, **p<0.01, ****p<0.0001; two-way ANOVA with Tukey’s *post hoc* HSD test for multiple comparisons.

Five (± 2) days prior to the end of the treatment period, mice were fasted overnight, before blood was sampled from the lateral tail vein for determination of fasting blood glucose and clinical chemistry analyses (**Suppl. Methods**). At 4-, 8- or 12-months of age (±2 days) mice were terminally anaesthetized *via* an intraperitoneal injection of sodium pentobarbital (500 mg.kg^-1^; Euthatal Solution, Merial Animal Health Ltd., Bracknell, UK). A terminal (fed-state) blood sample was collected *via* cardiac puncture, and the plasma separated (4000 x *g*, 10 min, 4°C; K_3_EDTA anticoagulant) and snap frozen for later analyses. The heart, liver, soleus and gastrocnemius muscles were rapidly collected (**Suppl. Methods**). Body composition was determined by dual energy x-ray absorptiometry (DEXA; **Suppl. Methods**; Lunar PIXImus II, GE Medical Systems Ltd., Chalfont St. Giles, UK). The left tibia was dissected and boiled for >5 hr to remove all soft tissue, then precisely measured from the tibial plateau to the tip of the medial malleolus.

### Metabolic Cages

For the 12–month cohort, mice underwent a 48–hour metabolic cage (Promethion®, Sable Systems, Germany) protocol at 10 months (± 6 days) of age (**Suppl. Methods**). Data were collected with a 1 Hz sampling rate and acquired and coordinated by the IM3 Interface Module (v20.0.6, Sable Systems), with real-time measurement of respiratory exchange ratio (RER) and calculation of energy expenditure (EE, *via* the Weir equation; [43]). Data were processed using ExpeData (v1.9.27, Sable Systems) with Macro Interpreter (v2.32 Sable Systems) to process raw data into 5 min bins. Metabolic cage data was corrected to body mass, lean mass or fat mass covariates, as appropriate (44), determined by time domain nuclear magnetic resonance (TD-NMR; **Suppl. Methods**). Data were analyzed using the CalR framework (45), adapted for a 2x2 factorial design, in R (46).

### Blood Analyses

Insulin, FFAs, TAGs, total cholesterol, and HDL-cholesterol were measured in fasted blood samples by the Core Biochemical Assay Laboratory (Cambridge University Hospitals, NHS Foundation Trust, Cambridge, UK). LDL-cholesterol was calculated using the Friedewald equation (47). Fasted glucose and insulin measurements were used to calculate the homeostatic model assessment (48) as a measurement of insulin resistance (HOMA-IR) adapted for murine models, as previously described (49).

Plasma nitrate and nitrite concentrations were determined in the fed state using a dedicated high-performance liquid chromatography analysis system (Eicom ENO-30, coupled to an AS-700 INSIGHT autosampler; Amuza Inc.), following methanol deproteinization as previously described (50). Plasma concentrations of the liver enzymes alanine transaminase (ALT) and aspartate transaminase (AST) were quantified *via* commercial ELISAs (#ab28282 and #ab263882, Abcam, Cambridge, UK). The antioxidant capacity of plasma was determined *via* the ferric reducing ability of plasma (FRAP) assay, as previously described (51). Lipid peroxidation was used as a marker of oxidative stress and quantified in plasma by measurement of thiobarbituric acid reactive substances (TBARS), as described (51).

### Intestinal Morphology

For investigation of intestinal morphology, an additional cohort of male C57Bl/6J mice (aged 8 weeks, n=7) was purchased from the same commercial breeder and randomized to receive either standard laboratory chow (RM3(E)) or HFHS diet (TD.88137). After 14 weeks, mice were terminally anaesthetized, and the gastrointestinal tract dissected from the pyloric sphincter to the rectum. After the caecum was removed, the intestinal tract was cut into 4 sections (colon, plus proximal, mid, and distal small intestine approximately equivalent to the duodenum, jejunum and ileum), and prepared for histological processing, as previously described (52).

### Hepatic Lipid Content

Hepatic lipid content was quantified using an optimized Folch extraction protocol for quantification of liver-fat (53).

### High-Resolution Respirometry

High-resolution respirometry was performed using an Oxygraph-2k (Oroboros Instruments, Innsbruck, Austria) on saponin-permeabilized muscle fiber bundles from the cardiac apex, soleus and gastrocnemius muscles and on liver left lateral lobe (LLL) homogenate, prepared as previously described (54, 55). A substrate-uncoupler-inhibitor titration was carried out to assess mitochondrial capacity as described in **Table 1**.

**Table 1.**
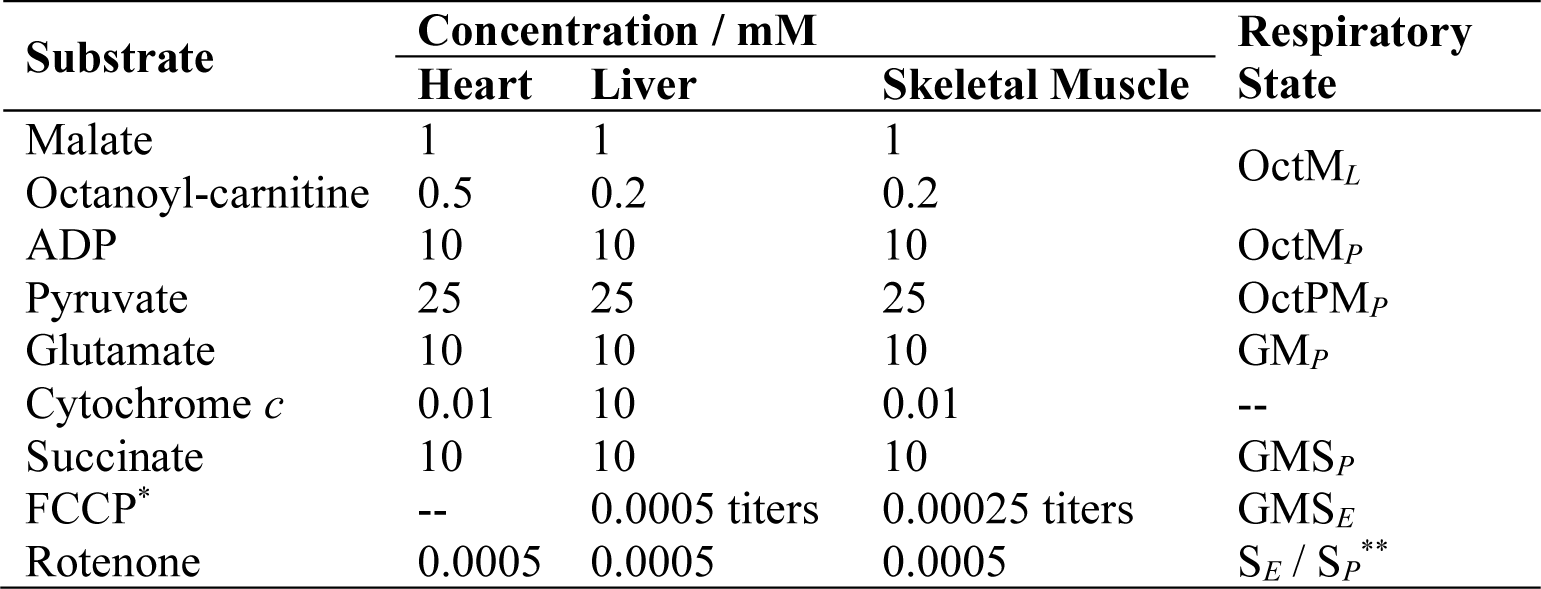
Substrate-uncoupler-inhibitor titration used to investigate mitochondrial capacity in permeabilized heart fibers, skeletal muscle fibers, and liver homogenate. Two types of skeletal muscle (gastrocnemius and soleus) were assayed, using the same titration protocol. Cytochrome *c* was added as a quality control to check outer mitochondrial membrane integrity. *Carbonyl cyanide-*p*-trufluoromethoxyphenylhydrazone (FCCP) was titrated in the indicated quantities until the maximum rate was reached. No FCCP was added in the heart as there was no reserve capacity measurable above maximal OXPHOS stimulated following saturation of the electron transport system (ETS) with glutamate and succinate. **In the heart assay, due to the absence of the uncoupler FCCP, addition of the complex I inhibitor rotenone led to measurement of OXPHOS supported by succinate via ETS complex II alone (state S*_P_*), whilst in liver and muscle the uncoupling of the ETS from ATP synthase meant the oxygen consumption rate measured reflected capacity of ETS complex II (state S*_E_*).

Flux control ratios were calculated to determine the proportion of maximal oxidative phosphorylation (OXPHOS) that could be supported by the F-pathway via β-oxidation (FCR_F_; [1]) and by the N-pathway via complex I (FCR_N_; [2]), as well as the relative capacity to support OXPHOS using octanoyl-carnitine and malate as substrates compared with pyruvate and malate (FCR_FA/P_; [3]). Mitochondrial OXPHOS coupling efficiency (OCE; [4]) was also calculated as a measure of the proportion of OXPHOS not limited by LEAK-state respiration.

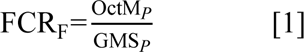

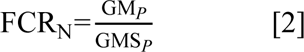

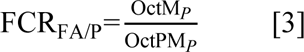

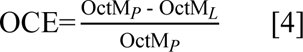

### Liquid Chromatography-Mass Spectrometry Based Metabolite Analyses

Aqueous and organic metabolites were extracted from ∼25 mg liver LLL or cardiac left ventricle using a modified Bligh and Dyer method (56), as previously described (57), dried under nitrogen and stored at -80°C until analysis.

For analysis of cardiac glycogenesis intermediates, half the aqueous fraction was reconstituted for hydrophobic interaction liquid chromatography (HILIC), which was performed as previously described (58) using Vanquish UHPLC^+^ series coupled to a TSQ Quantiva Triple quadrupole Mass Spectrometer (both Thermo Scientific). Identification of detected metabolites was performed using targeted analysis of parent/daughter ion *m/z* and retention times from chemical standards: glucose: +179.000→+89.000, 2.6 min; glucose-6-phosphate: +259.000→+97.077, 4.0 min; glucose-1-phosphate: +259.000→+97.077, 3.9 min; UDP-glucose: +565.078→+323.000, 3.88 min.

Analysis of acyl-carnitines in liver and heart, was carried out using a reconstituted mix of half of each of the aqueous and organic phases, as previously described (55). Acylcarnitine analysis was carried out on the same instrumentation as HILIC analysis, with data collection and processing in both cases carried out using Xcalibur software (v.2.2, Thermo Scientific).

Open-profiling lipidomics was carried out as previously described (57) with lipid data collected using the Fourier transform mass spectrometer analyzer set in profile mode with a resolution of 60,000 (Ultimate 3000 UHPLC system) coupled to an LTQ Orbitrap Elite Mass Spectrometer [both Thermo Fisher Scientific]). A full scan was performed across an *m/z* range of 110–2000. Lipid metabolites were processed as previously described (57) with peaks identified by XCMS software (59) based on an approximate chromatographic peak detection (FWHM) of 5 seconds and a signal to noise threshold ratio of 5. Peaks were annotated by accurate mass, using an automated in–house R script and by comparison to the LipidMaps database (60).

All data were normalized to the intensity of appropriate internal standards and to sample protein concentration (Pierce BCA Assay, Thermo Scientific) to account for differences in sample size.

### Reverse Transcription-Quantitative PCR

Total RNA was extracted from ∼30 mg liver LLL or ∼25 mg cardiac left ventricle using an RNeasy Fibrous Tissue Mini kit (QIAGEN Ltd., Manchester, UK) according to the manufacturer’s instructions. RNA (1 μg, quantified *via* Nanodrop-2000 spectrophotometer) was transcribed to cDNA using the QuantiTect Reverse Transcription kit (QIAGEN) and stored at -20°C prior to analysis. Real time qPCR was carried out using a LightCycler® 480 System (Roche) fitted with a 384-well block, using the QuantiNova SYBR Green PCR kit (QIAGEN) and QuantiTect primer assays (QIAGEN; **Suppl. Table 1**). Expression was determined in triplicate from each biological replicate. Fold-change in gene expression was determined using the relative standard curve method, with expression of all genes normalized to the geometric mean of three housekeeping genes: *Rn18s*, *Srsf4* and *Hmbs* in liver, and *Actb*, *Ywhaz* and *Rn18s* in heart.

### Histology

Sections were fixed in PFA (cardiac mid-sections) or formalin (LLL and intestinal sections) and embedded into paraffin blocks using standard laboratory methods. For LLL sections, tissue was processed at the Histopathology Core (MRC Metabolic Diseases Unit, Cambridge). All tissue was sectioned at 7 μm using an RM2235 Microtome (Leica Biosystems).

Liver and intestinal morphology was determined via hematoxylin (#ab220365, Abcam) and eosin staining (H&E) using standard laboratory protocols. Cardiac and hepatic fibrosis was determined by staining for collagen using picrosirius red (PSR, **Suppl. Methods**) whilst cardiac glycogen was stained using Periodic Acid Schiff (PAS, **Suppl. Methods**). Following staining, slides were dehydrated in ethanol, cleared in xylene and mounted using DPX. Fix-frozen right lateral lobe was used to histologically assess hepatic steatosis following cryosectioning at 10 μm; neutral lipid was stained with oil red O (ORO) and slides mounted in glycerin gelatin (**Suppl. Methods**).

Slides were imaged with a NanoZoomer 2.0-RS with images collect using NDP.view2 (v2.9.25, Hamamatsu Photonics, Japan). At least 3 independent sections per biological replicate were examined and analyzed. To determine the level of hepatic pathology, a MASLD activity score (MAS) was generated for each section using sections blinded for experimental condition (**Suppl. Table 2**; [61]). Quantification of villus length was carried out manually in NDP.view2, with 10 intact villi measured per quadrant per intestinal section (62). Quantification of PSR and PAS in cardiac sections were determined following color deconvolution in ImageJ (63). PAS staining was corrected for that in control sections treated with α-amylase to account for any non-glycogen PAS staining.

### Statistics

All data are presented as mean ± standard error, unless otherwise stated, with differences accepted as statistically significant when p ≤ 0.05. All univariate statistical tests were carried out in R (45) with graphs produced in Prism (v.10.0.3, GraphPad Software LLC., Boston, MA, USA). Multivariate statistical analysis was carried out using SIMCA-17 (Umetrics, Sartorius Stedim Data Analytics AB, Umeå, Sweden). Unless otherwise stated, statistical analysis was carried out in line with two *a priori* questions outlined before commencing the study: (a) at each time point, is there an effect of diet or nitrate supplementation? (b) is there an effect of diet or nitrate supplementation on age-related and/or disease progression? For each question, 2-way ANOVAs were carried out on the relevant experimental cohorts as appropriate, with correction for false discovery rates as required (**Suppl. Fig. 1**). When significant interactions were found, Tukey’s *post hoc* honest significance difference (HSD) test was used to consider relevant interactions (**Suppl. Fig. 1**).

## Results

### Whole-body metabolic phenotype

Mice fed a HFHS diet developed obesity that persisted throughout the 44 weeks of the study (**Fig. 1B**). After 3 weeks on the respective diets, HFHS-fed mice were 4.6% heavier than chow-fed mice (p<0.01) and by 12-months of age, HFHS-fed mice were 37.6% heavier than their counterparts on the control chow diet (p<0.0001; **Fig. 1B**). There was no effect of nitrate supplementation on body mass (**Fig. 1B**).

Food intake remained constant throughout the study, with chow-fed mice consuming 4.0 ± 0.03 g.mouse^-1^.day^-1^ and HFHS-fed mice eating 3.1 ± 0.05 g.mouse^-1^.day^-1^. However, as the HFHS-diet was more calorie-dense, there was no significant difference in energy intake between the experimental groups throughout the study (**Table 2**). HFHS-fed mice drank 27.9% less water per day (p<0.01; **Table 2**) compared with chow-fed controls, likely due to the higher sodium content of latter (0.3% in HFHS *vs.* 10% in chow diet). Despite this, mice receiving drinking water supplemented with 1 mM NaNO_3_ had a 5-fold higher intake of nitrate compared with NaCl-supplemented controls (p<0.0001; **Table 2**). This resulted in a plasma nitrate concentration that was 1.9-fold higher than in NaCl-supplemented controls at 4 and 8-months of age (p<0.05; **Table 3**). Circulating nitrite concentrations were significantly elevated only in 4-month-old mice fed standard rodent chow (**Table 3**). At 12-months there were no differences in plasma nitrate levels between any of the experimental groups (**Table 3**).

**Table 2.** Food, water and NO_3_^-^ intake across the experimental groups. Values were calculated based on individual cage measurements (n=3-4 mice/cage; 6 cages/group). Nitrate intake represents combined intake from supplemented water and food (using values provided from the diet suppliers). Data show mean ± SEM. **p<0.01, ****p<0.0001, compared with chow-fed mice receiving the same water treatment; ^††††^p<0.0001 compared with chloride-supplemented mice receiving the same diet; two-way ANOVA with Tukey’s HSD post hoc test for multiple comparisons.

|  | **Food Intake /**<br>kcal.mouse<sup>-1</sup>.day<sup>-1</sup> | **Water Intake /**<br>ml.mouse<sup>-1</sup>.day<sup>-1</sup> | **NO<sub>3</sub><sup>-</sup> Intake /**<br>mg.kg<sup>-1</sup>.day<sup>-1</sup> |
| --- | --- | --- | --- |
| Chow / Chloride | 14.5 ± 0.1 | 5.8 ± 0.1 | 1.4 ± 0.0 |
| Chow / Nitrate | 14.1 ± 0.1 | 5.3 ± 0.1 | 12.4 ± 0.2<sup>††††</sup> |
| HFHS / Chloride | 13.8 ± 0.2 | 4.1 ± 0.1<sup>**</sup> | 1.9 ± 0.1 |
| HFHS / Nitrate | 14.9 ± 0.2 | 3.9 ± 0.1<sup>**</sup> | 8.2 ± 0.2<sup>****††††</sup> |

**Table 3.**
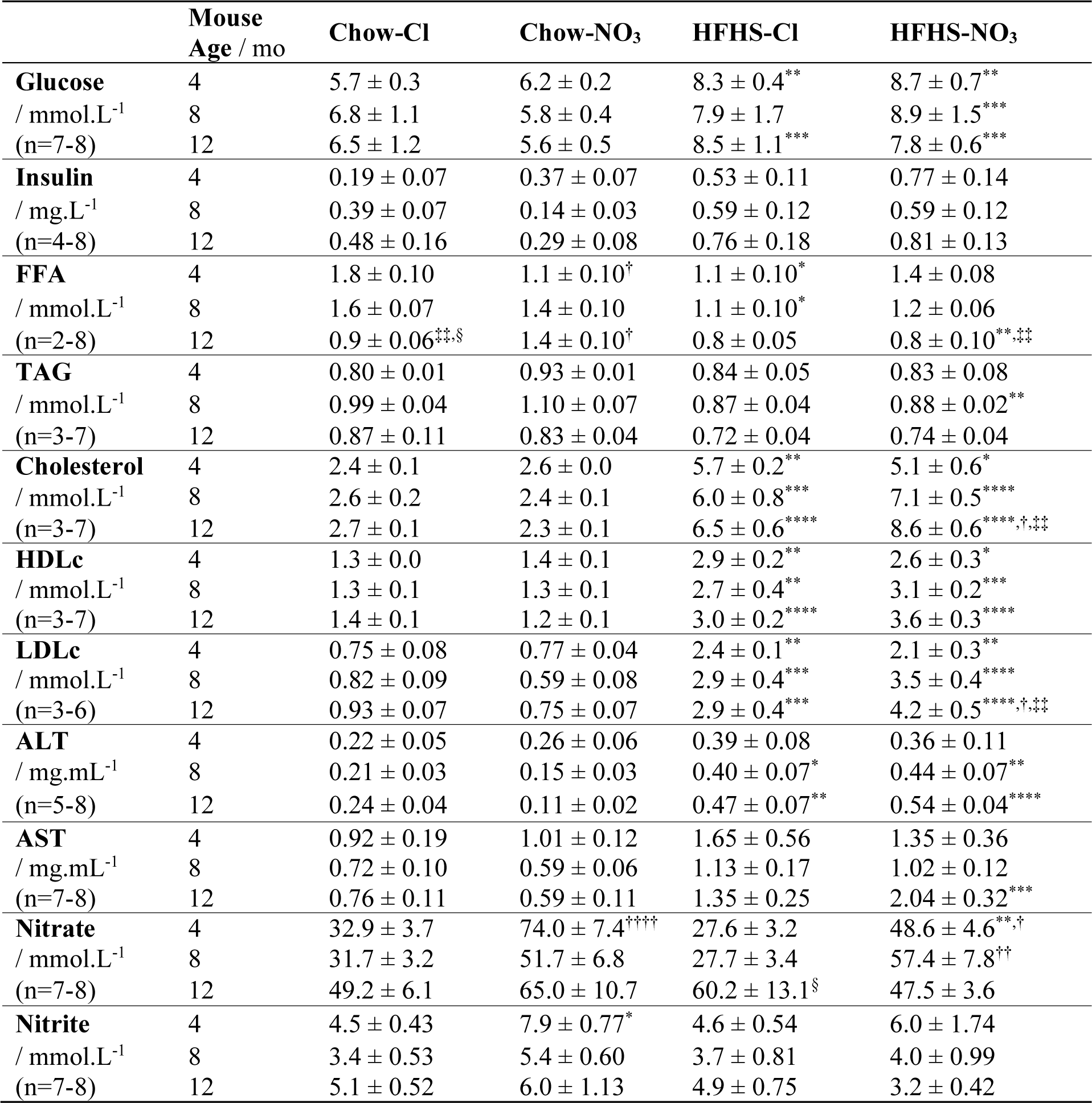
Clinical biochemistry measurements and other plasma parameters. All clinical biochemistry measurements were assayed from fasted plasma. Nitrate and nitrite concentrations, and liver enzymes were measured from terminal blood samples (fed state). Data show mean ± SEM with N numbers assayed indicated. FFA = free fatty acids; TAG = triacylglycerols; HDLc = high density lipoprotein cholesterol; LDLc = low density lipoprotein cholesterol; ALT = alanine transaminase; AST = aspartate transaminase. *p<0.05, **p<0.01, ***p<0.001, ****p<0.0001, compared with chow-fed mice receiving the same water treatment; ^†^p<0.05, ^††^p<0.01, ^††††^p<0.0001 compared with chloride-supplemented mice receiving the same diet; ^‡‡^p<0.01 compared with 4-month-old mice within the same treatment group; ^§^p<0.05 compared with 8-month-old mice within the same treatment group; all statistics represent two-way ANOVA with Tukey’s HSD post hoc test for multiple comparisons.

The greater body mass observed in HFHS-fed mice compared with chow-fed mice was reflected in a greater body fat percentage, and correspondingly, a lower lean body mass percentage (p<0.0001; **Fig. 1C, D**). Body fat was ∼2-2.5-fold greater, whilst lean mass was approximately one third less in mice fed a HFHS diet, compared with chow-fed controls. In the 8-month cohort, HFHS-fed mice also receiving NaNO_3_ had 31.3% more body fat and 13.3% less lean mass compared with chloride-supplemented counterparts on the HFHS diet (p<0.05), although this exacerbation of obesity in nitrate-fed mice was not seen at 12-months (**Fig. 1C, D**).

Mice fed a HFHS diet had a fasted blood glucose concentration that was 36.7% higher than that of chow-fed mice across all ages assessed (p<0.01), with no difference in fasted plasma insulin concentration (**Table 3**). Accordingly, HFHS-fed mice had ∼2-fold higher HOMA-IR scores indicating evidence of insulin resistance, which reached statistical significance in mice supplemented with NO_3_^-^ from 8-months of age compared with their chow-fed counterparts (p<0.05; **Fig. 1E**). Total, HDL and LDL cholesterol were elevated in mice fed a HFHS diet across all ages compared with chow-fed controls (2.6-fold, 2.1-fold and 3.7-fold higher respectively; p<0.05; **Table 3**). Total and LDL cholesterol were also elevated in the 12-month HFHS-NO_3_ cohort, compared with both the same group at 4-months of age (69.1% total, 2.1-fold LDL; p<0.01; **Table 3**), and with the 12-month HFHS-Cl group (30.9% total, 1.5-fold LDL; p<0.05; **Table 3**), suggesting an exacerbation of dyslipidemia with longer-term nitrate supplementation. Cholesterol ratio (total/HDL cholesterol, [64]) was also elevated in HFHS-NO_3_ mice at 8-months and 12-months of age, compared with chow-fed controls (23.1%; p<0.05; **Fig. 1F**) and raised at both of these time-points in comparison with 4-month-old HFHS-NO_3_-fed mice (20.2%, p<0.05; **Fig. 1F**). There were, however, few clear differences in the plasma free fatty acids or triacylglycerol (TAG) concentrations as a result of age, diet or nitrate treatment (**Table 3**).

We also detected elevations in plasma levels of the liver enzymes ALT and AST indicative of liver damage (65). In chloride-supplemented HFHS-fed mice, plasma ALT concentration was 92.4% higher at 8-months (p<0.05) and 96.4% higher at 12-months (p<0.01) in comparison with chow-fed controls, whilst in those receiving nitrate, plasma ALT concentration was 2.9-fold greater at 8-months (p<0.01) and 4.8-fold greater at 12-months (p<0.0001) compared with chow-matched controls (**Table 3**). Plasma AST concentration was also elevated in HFHS-fed mice: in the 8-month cohort, plasma AST was 64.3% higher than in chow fed mice (p<0.01) whilst at 12-months, HFHS-NO_3_ fed mice showed a plasma AST concentration 3.5-fold higher than chow-NO_3_ mice (p<0.001; **Table 3**).

Quantification of TBARS in plasma showed that, at every time point, lipid peroxidation was higher in HFHS-fed mice compared with chow-fed controls (2.6-fold; p<0.05; **Fig 1G**), with no effect of nitrate supplementation. This increase in lipid peroxidation was not associated with any differences in plasma antioxidant capacity, as shown by similar FRAP concentrations across all cohorts (**Fig. 1H**).

Taken together, this data does not support a protective effect of nitrate supplementation for metabolic health during diet-induced obesity.

### Whole-body energy balance

Next, we sought to determine whether HFHS-feeding or nitrate administration altered whole body energy balance, as has been previously reported (66, 67). We used indirect calorimetry and activity measurements to investigate energy utilization in the 12-month cohort whilst mice were 10 months of age (**Fig. 2A-E**).

**Figure 2.**
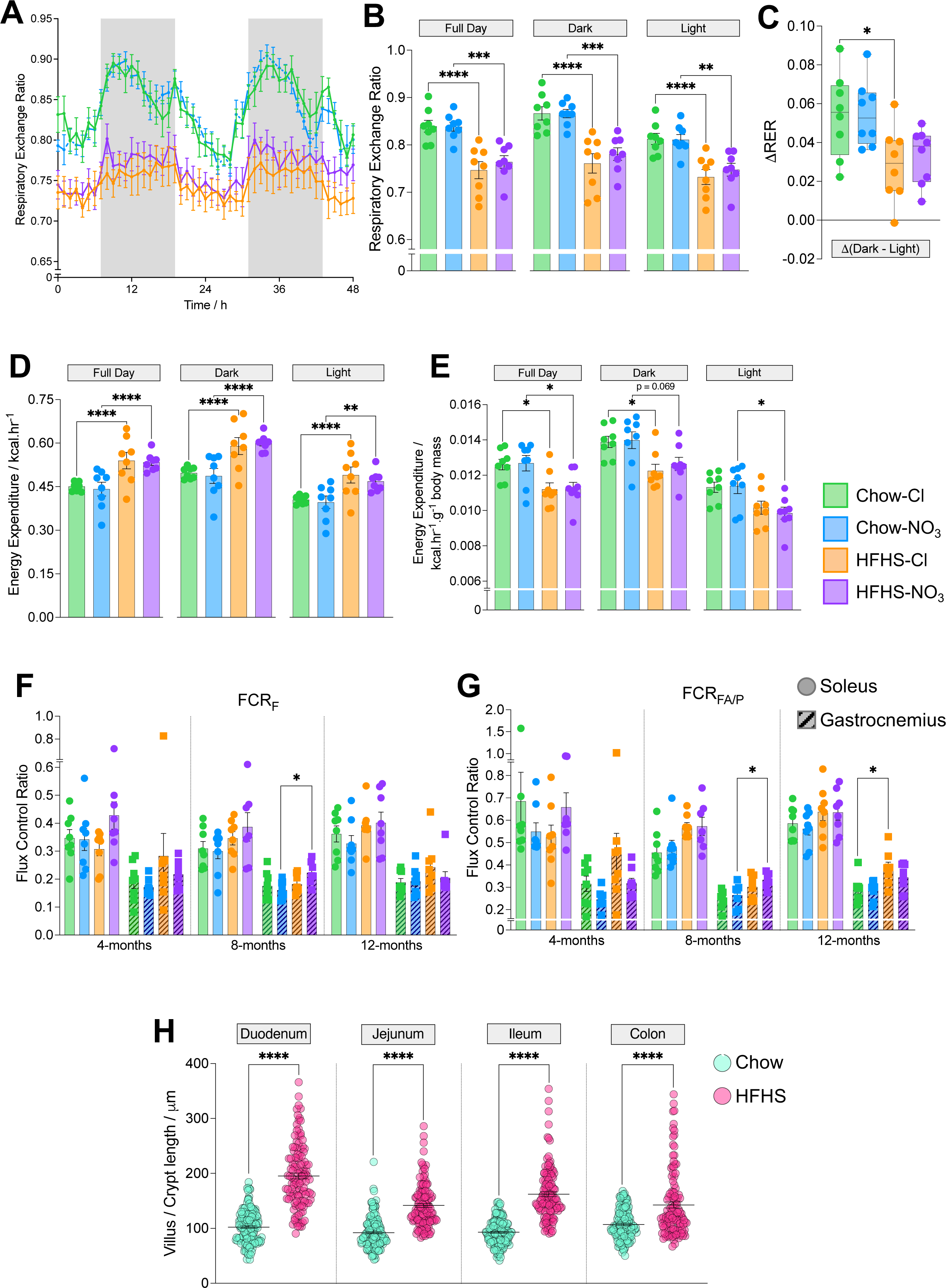
Whole body energy expenditure and skeletal muscle mitochondrial metabolism differences do not account for development of obesity in high-fat high-sucrose fed mice. (A) Changes in respiratory exchange ratio, determined from *V*O_2_ and *V*CO_2_. Shaded areas represent the dark period. (B) Average respiratory exchange ratio during the dark period, light period and across a full day, determined from *V*O_2_ and *V*CO_2_ in 10-month-old mice during the 48-hour metabolic cage protocol. (C) Difference in average respiratory exchange ratio between the active dark period, and the quiet light period. (D) Energy expenditure across the dark period, light period and full day. (E) Mass normalised energy expenditure (normalised to total body mass) across the dark period, light period and full day. For (B)-(E) N=8/group; *p<0.05, **p<0.01, ***p<0.0001, ****p<0.0001; analyzed according to the CalR framework (44) adapted for a 2x2 design. (F) Contribution of the F-pathway *via* β-oxidation to maximal OXPHOS in saponin-permeabilized skeletal muscle fibers. (G) Capacity for OXPHOS supported by octanoyl-carnitine and malate relative to pyruvate and malate, in saponin-permeabilized skeletal muscle fibers. For (F)-(G) N=7-8/group; *p<0.05; two-way ANOVA with Tukey’s post hoc HSD test for multiple comparisons. (H) Villus length (or colonic crypt depth) across regions of the intestine measured in histological sections (**Suppl.** Fig. 2D) from mice fed chow or HFHS-diet for 8-weeks. N=3-4/group; ****p<0.0001; unpaired two-tailed t-test with Welch’s correction. All data represent mean ± SEM.

Throughout the 48-hour metabolic cage protocol, chow-fed mice had a higher respiratory exchange ratio (RER) than HFHS-fed mice (0.84 ± 0.005 *vs.* 0.76 ± 0.002; p<0.001) that was independent of nitrate supplementation (**Fig. 2A, B**), indicating that HFHS-fed mice were more reliant on fat-oxidation than their chow-fed counterparts (68). Chow-fed mice also showed a greater difference in RER between the peaks and troughs recorded during light and dark periods compared with HFHS-fed mice (**Fig. 2A-C**), indicating greater metabolic flexibility to switch between the fed and fasted states, and this reached statistical significance in NaCl-supplemented mice (47.0% lower in HFHS-Cl mice, p<0.05; **Fig. 2C**).

HFHS-fed mice had a total energy expenditure (EE) that was 20.3% greater than mice maintained on the chow diet (p<0.0001; **Fig. 2D**), with no effect of nitrate supplementation. When normalized to body mass, however, EE was 11.2% lower in HFHS-fed mice, compared with chow-fed mice (p<0.05; **Fig. 2E**). Again, there was no apparent effect of nitrate supplementation, although mass-corrected EE was 13.5% lower in the inactive light period in HFHS-NO_3_ mice compared with chow-fed controls (p<0.05; **Fig. 2E**). Regression analysis demonstrated that EE varied consistently with total body mass across all experimental groups (**Suppl. Fig. 2A**).

Given the relationship between EE and body mass, we investigated how the HFHS-diet might exert an obesogenic effect despite similar caloric intake and EE compared with chow-fed controls. First, we investigated mitochondrial respiratory capacity in the skeletal muscle, since this is the largest insulin-sensitive tissue in the body (69) and largely dominates the measurement of whole-body RER (70). We found very little difference in mitochondrial respiration between cohorts in either soleus (oxidative type-I muscle) or gastrocnemius (a more glycolytic type-II muscle, [71]; **Suppl. Fig 2B, C**), with similar OXPHOS coupling efficiencies across treatment groups, within given muscle types (**Suppl. Fig. 2D**).

However, contribution to maximal OXPHOS through the F-pathway *via* β-oxidation (FCR_F_) was 38.8% higher in gastrocnemius from 8-month-old HFHS-NO_3_ mice compared with chow-fed controls (p<0.05; **Fig. 2F**) and the relative capacity for fatty acid oxidation (FCR_FA/P_) was also higher in gastrocnemius from HFHS-fed mice compared with chow-fed controls (26.0% higher in 8-month-old HFHS-NO_3_ mice; 32.6% higher in 12-month HFHS-Cl mice; p<0.05; **Fig. 2G**). These differences were not apparent in soleus (**Fig. 2F, G**).

These findings support a shift in mitochondrial substrate preference in the gastrocnemius of HFHS-fed mice, but did not indicate a reduction in overall muscle oxidative capacity or altered mitochondrial efficiency.

Next, we investigated whether the HFHS diet resulted in similar changes in intestinal morphology to those previously reported to result from dietary fructose (62). Accordingly, a cohort of mice fed the HFHS diet for 8-weeks showed longer villi across the small intestine and deeper colonic crypts than chow-fed controls (62.8%, p<0.0001; **Fig. 2H**, **Suppl. Fig. 2E**). This suggests that obesity in the HFHS-fed mice resulted from increased nutrient absorption rather than increased intake or reduced expenditure.

### Cardiac metabolism and fibrosis

As ∼65% of diabetes-related mortality arises from cardiovascular diseases (72), with alterations in myocardial metabolism considered central to the etiology (73), we sought to investigate whether inorganic nitrate supplementation altered cardiac metabolic function in HFHS-fed mice. There was no difference in either heart mass (**Suppl. Fig. 3A**) or heart mass normalized to tibia length (a proxy for body size [74]) as a result of age, diet or nitrate supplementation (**Fig. 3A**).

**Figure 3.**
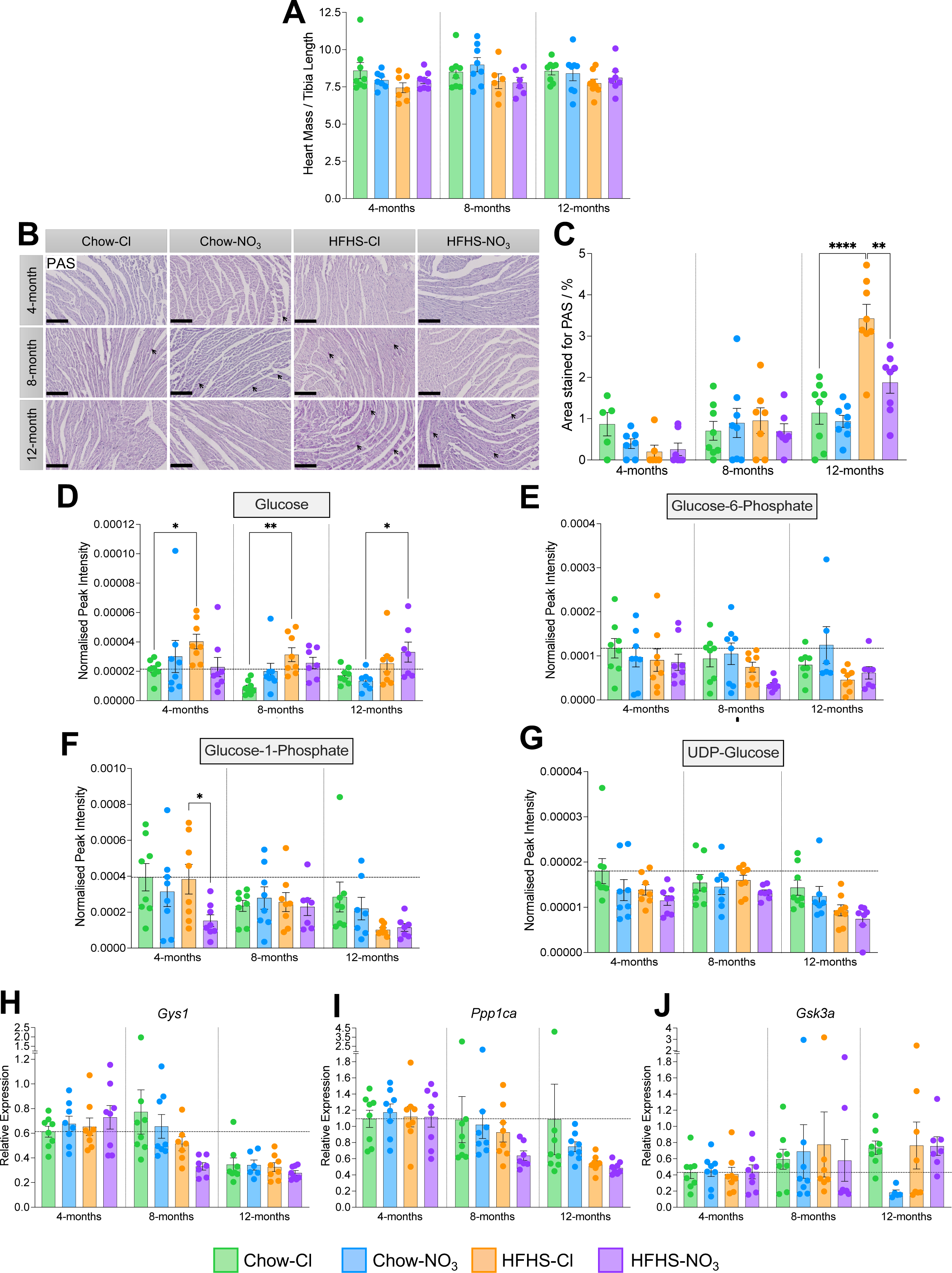
Inorganic nitrate supplementation prevents glycogen accumulation in HFHS-fed mouse heart. (A) Heart mass, normalised to tibial length, as a proxy for body size. (B) Representative cardiac midsections stained with Periodic Acid Schiff (PAS) stain at 20x magnification. Scale bars represent 150 mm. Glycogen deposits are stained magenta and marked with arrows (maximum 3 per slide shown). (C) Quantification of area staining positive for PAS. Percentages were corrected for the staining present in control sides pre-treated with a-amylase, thereby excluding any staining resulting from other glycans and mucins. (D) Relative cardiac concentration of glucose, determined by hydrophobic interaction liquid chromatography (HILIC)-mass spectrometry (MS). (E) Relative cardiac concentration of glucose-6-phosphate, determined by HILIC-MS. (F) Relative cardiac concentration of glucose-1-phosphate, determined by HILIC-MS. (G) Relative cardiac concentration of UDP-glucose, determined by HILIC-MS. (H) Relative cardiac expression of *Gys1*, determined by RT-qPCR. (I) Relative cardiac expression of *Ppp1ca*, determined by RT-qPCR. (J) Relative cardiac expression of *Gsk3a*, determined by RT-qPCR. Data represent mean ± SEM; N=6-8 / group; data in (D)-(G) are normalised to an appropriate internal standard and sample protein concentration; For (H)-(J) horizontal line indicates relative expression in 4-month-chow/chloride mice. *p<0.05, **p<0.01, ***p<0.001, ****p<0.0001; two-way ANOVA with Tukey’s *post hoc* HSD test for multiple comparisons.

Cardiac glycogen content was assessed in sections of cardiac midsection using PAS staining (**Fig. 3B**, **C**). When corrected for staining in the presence of a-amylase, the hearts of 12-month-old HFHS-fed mice showed 3.0-fold higher PAS staining than those of age-matched chow-fed controls (p<0.0001), indicative of increased glycogen content (**Fig. 3C**). The area staining positively for PAS was also 82.5% higher in the HFHS-Cl cohort than the HFHS-NO_3_ cohort at 12-months of age (p<0.01, **Fig. 3C**), indicating that glycogen accumulation was prevented when mice were supplemented with nitrate.

Using HILIC-mass spectrometry, we quantified metabolites involved in glycogen synthesis in the left ventricle (**Fig. 3D-G**). HFHS-fed mice had 59.6% higher cardiac glucose levels compared with chow-fed controls across all cohorts (**Fig. 3D**), which may be due to the high sucrose content of the diet. HFHS-Cl-fed mice had 87.5% higher cardiac glucose at 4-months (p<0.05) and 3.5-fold higher cardiac glucose at 8-months (p<0.01) compared with chow-Cl mice, whilst HFHS-NO_3_-fed mice showed a 2.5-fold higher cardiac glucose than chow-fed controls at 12-months of age (p<0.05, **Fig. 3D**). Cardiac glucose-6-phosphate showed no difference across age, diet or nitrate supplementation (**Fig. 3E**), however glucose-1-phosphate levels were 60.3% lower in hearts from HFHS-NO_3_ mice compared with HFHS-Cl mice at 4-months of age (p<0.05; **Fig. 3F**). Glucose-1-phosphate levels in HFHS-Cl fed mice, however, showed decreasing levels with age (**Fig. 3F**), being 33.2% lower in 8-month-old mice compared to 4-month-old mice, and 73.5% lower in 12-month-old mice fed HFHS-Cl (p<0.05; **Fig. 3F**). There was no difference in cardiac levels of UDP-glucose as a result of diet or nitrate supplementation (**Fig. 3G**). However, compared with 8-month-old mice, hearts from 12-month-old mice fed a HFHS diet showed a lower level of UDP-glucose, with detected levels being 43.0% lower in the hearts of HFHS-fed mice (p<0.05) at 12-months compared with those at 8-months of age (**Fig. 3G**).

The expression of genes involved in glycogen synthesis was unaffected by diet or nitrate supplementation (**Fig. 3H-J**), however, both glycogen synthase 1 (*Gys1*) and *Ppp1ca* (one of the catalytic subunits of protein phosphatase 1) showed lower expression in the hearts of older HFHS-fed mice. In HFHS-fed mice receiving NaCl, *Gys1* expression was 50.2% lower (p<0.05; **Fig. 3H**) and *Ppp1ca* was 51.8% lower (p<0.01; **Fig. 3I**) at 12-months than at 4-months. However, in HFHS-NO_3_-fed mice, lower expression was detectable in the 8-month cohort, with *Gys1* expression 54.5% lower at 8-months (p<0.001) and 61.8% lower expression at 12-months (p<0.0001) compared with hearts from 4-month-old mice (**Fig. 3H**), and *Ppp1ca* expression 42.8% lower at 8-months (p<0.05) and 56.5% lower at 12-months (p<0.001) compared with that at 4-months (**Fig. 3I**). There were no differences in cardiac expression of *Gsk3a* as a result of age or disease progression (**Fig. 3J**).

Taken together, these findings suggest that nitrate supplementation minimizes glycogen accumulation as a result of long-term HFHS feeding. Glycogen accumulation, which can occur in the diabetic heart (75) has been linked to the cardiometabolic stress response (76).

We next investigated myocardial fat metabolism, given that fatty acids are the predominant source of ATP in the healthy heart (77) and alterations in substrate usage are canonical markers of both the diabetic and failing heart (73, 78). Total TAG content of the left ventricle (as determined by LC-MS peak intensity) was 61.1% higher at 4-months (p<0.05) and 2.4-fold higher at 8-months (p<0.01) in HFHS-fed mice compared with chow-fed controls (**Fig. 4A**). However, in nitrate-supplemented HFHS-fed mice, cardiac TAGs were not greater than in chow-fed mice (**Fig. 4A**). Similarly, total cardiac diacylglycerol (DAG) was 53.1% higher in HFHS-Cl mice at 4-months (p<0.0001) and 79.8% higher at 8-months (p<0.0001) compared with chow-fed controls (**Fig. 4B**), whilst HFHS-NO_3_ fed mice showed no differences in cardiac DAG levels compared with chow-fed mice. However, chow-NO_3_-fed mice showed higher cardiac levels of DAGs at 4-months (+29.6%, p<0.05) and at 8-months (+31.3%, p<0.05) compared with aged-matched chow-Cl mice (**Fig. 4B**). Inorganic nitrate supplementation therefore limited lipid accumulation in the heart, as a result of HFHS-feeding.

**Figure 4.**
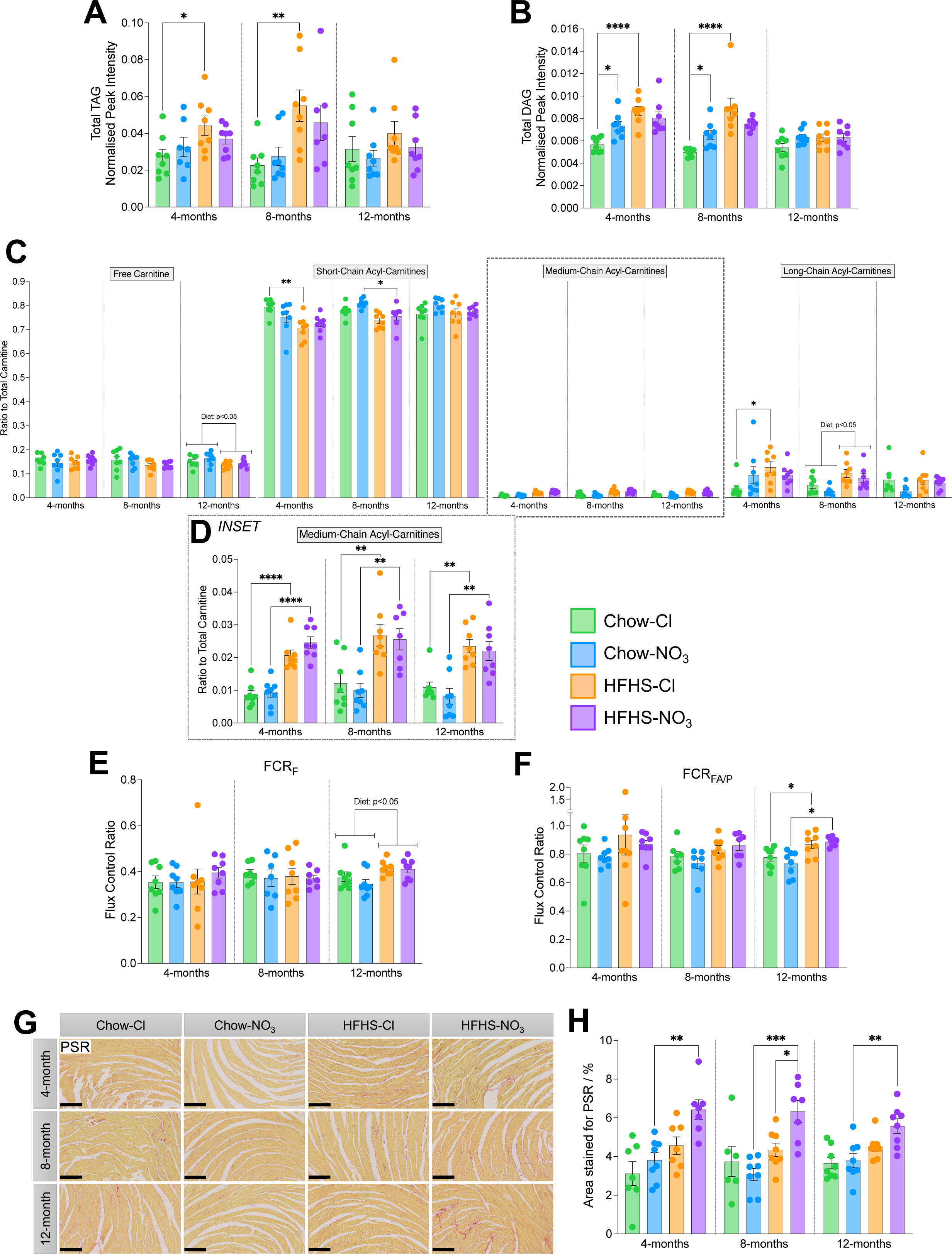
Inorganic nitrate supplementation offers minor benefits to cardiac fat metabolism in HFHS-fed mice, but worsens cardiac fibrosis. (A) Total cardiac triacylglycerol content, determined by liquid chromatography-mass spectrometry, measured as total peak area ratio, normalised to an appropriate internal standard and sample protein concentration. (B) Total cardiac diacylglycerol content, determined by liquid chromatography-mass spectrometry, measured as total peak area ratio, normalised to an appropriate internal standard and sample protein concentration. (C) Relative levels of free (no associated acyl chain), short-chain (C2-C5), medium-chain (C6-C12) and long-chain (≥C13) acyl-carnitines to the total carnitine pool as measured by liquid chromatography-mass spectrometry (LC-MS) from cardiac tissue. (D) Relative levels of medium-chain acyl-carnitines to the total carnitine pool (from (H)) shown as an expanded inset. (E) Contribution of the F-pathway *via* β-oxidation to maximal OXPHOS in saponin-permeabilized cardiac fibers. (F) Capacity for OXPHOS supported by octanoyl-carnitine and malate relative to pyruvate and malate, in saponin-permeabilized cardiac fibers. (G) Representative cardiac midsections stained with picrosirius red (PSR) at 10x magnification. Scale bars represent 250 mm. Collagen fibers are stained red, identifying regions of fibrosis. (H) Quantification of area staining position for picrosirius red, as an indicator of cardiac fibrosis. Data represent mean ± SEM; N=6-8 / group. *p<0.05, **p<0.01, ***p<0.001, ****p<0.0001; two-way ANOVA with Tukey’s *post hoc* HSD test for multiple comparisons.

Moreover, combined HFHS-NO_3_ feeding prevented a reduction in the total acyl-carnitine pool that was observed in hearts from mice fed HFHS-Cl at 4-months (-21.2%, p<0.05) and 12-months (- 35.2%, p<0.01) compared with chow-fed controls (**Suppl. Fig. 3B**). However, HFHS-feeding resulted in alterations to the composition of the cardiac acyl-carnitine pool, independent of nitrate supplementation (**Fig. 4C**, **D**; **Suppl. Fig. 3C**). There was a relative depletion in the contribution of short-chain (total carbons C2–C5) acyl-carnitines to the total cardiac carnitine pool in mice fed a HFHS diet (**Fig. 4C**), reaching significance in the 4-month-old HFHS-Cl mice (11.0% lower than chow-Cl hearts, p<0.01) and 8-month-old HFHS-NO_3_ mice (6.6% lower than chow-NO_3_ hearts, p<0.05), whilst long-chain acyl-carnitines (total carbons ³C13) showed a 3.3-fold higher contribution to the total cardiac carnitine pool in 4-month-old HFHS-Cl fed mice (p<0.05) and 2.3-fold higher in HFHS-fed mice, independent of nitrate supplementation at 8-months (p<0.05; **Fig. 4C**). Medium-chain acyl-carnitines (total carbons C6-C12) were higher in HFHS-fed mice, regardless of age or nitrate supplementation (p<0.01; **Fig. 4C**, **D**).

To investigate whether these changes in metabolites were reflected in mitochondrial capacity, we investigated cardiac mitochondrial respiratory function in saponin-permeabilized fiber bundles. We found no differences in mass-specific respiration in any state assayed, due to diet, nitrate nor age (**Suppl. Fig. 3D**). However, at 12 months, relative contribution of fatty acid oxidation (FCR_F_) was 13.9% higher in the hearts of HFHS-fed mice than those of chow-fed mice regardless of nitrate supplementation (p<0.01, **Fig. 4E**). The capacity for fatty acid oxidation relative to pyruvate oxidation (FCR_FA/P_) was also higher in the hearts of 12-month-old HFHS-fed mice: HFHS-Cl fed mouse hearts had 11.8% higher FCR_FA/P_ compared with chow-Cl mouse hearts (p<0.05), whilst HFHS-NO_3_ mice had a 20.9% higher cardiac FCR_FA/P_ than chow-NO_3_ mice (p<0.05, **Fig. 4F**).

We also investigated cardiac fibrosis histologically in cardiac midsections using picrosirius red (PSR) staining for collagen (**Fig. 4G**). Analysis of the area staining positive for collagen showed that, at all ages, HFHS-NO_3_ fed mice had a 71.7% higher cardiac collagen content than chow-fed controls (p<0.01, **Fig. 4H**). At 8-months, hearts from HFHS-NO_3_ mice also showed a collagen content that was 45.4% higher than that of hearts from HFHS-Cl mice (p<0.05). This trend of higher collagen content in HFHS-NO_3_ mouse hearts compared with HFHS-Cl mice was also seen at 4-months (40.9% higher; p = 0.0624) and 12-months (23.8% higher; p = 0.107; **Fig. 4H**).

In summary, inorganic nitrate supplementation exerted a metabolic benefit in the HFHS-fed mouse heart, through minimizing cardiac lipid accumulation and delaying glycogen accumulation, but did not affect mitochondrial respiratory capacity, and resulted in increased cardiac fibrosis from 4 months.

### Inorganic nitrate and MASLD progression

A common, silent comorbidity of obesity is MASLD (79). We therefore investigated the effect of inorganic nitrate supplementation on liver metabolism and MASLD progression in HFHS-fed mice.

At 12-months, mice fed a HFHS-diet had livers that were 1.8-fold larger (relative to body mass) than chow-fed controls (p<0.0001; **Fig. 5A**). Interestingly, HFHS-NO_3_ fed mice showed greater hepatomegaly than HFHS-Cl fed mice (total liver mass being 1% heavier relative to body weight, p<0.05; **Fig. 5A**). HFHS-NO_3_ mice also had a greater prevalence of tumors, with 50% of this group having visible tumors on the liver surface, compared to only one mouse in the HFHS-Cl group and none in chow-fed mice (χ^2^(df = 3, N = 32) = 10.2, p<0.05; **Fig. 5B**).

**Figure 5.**
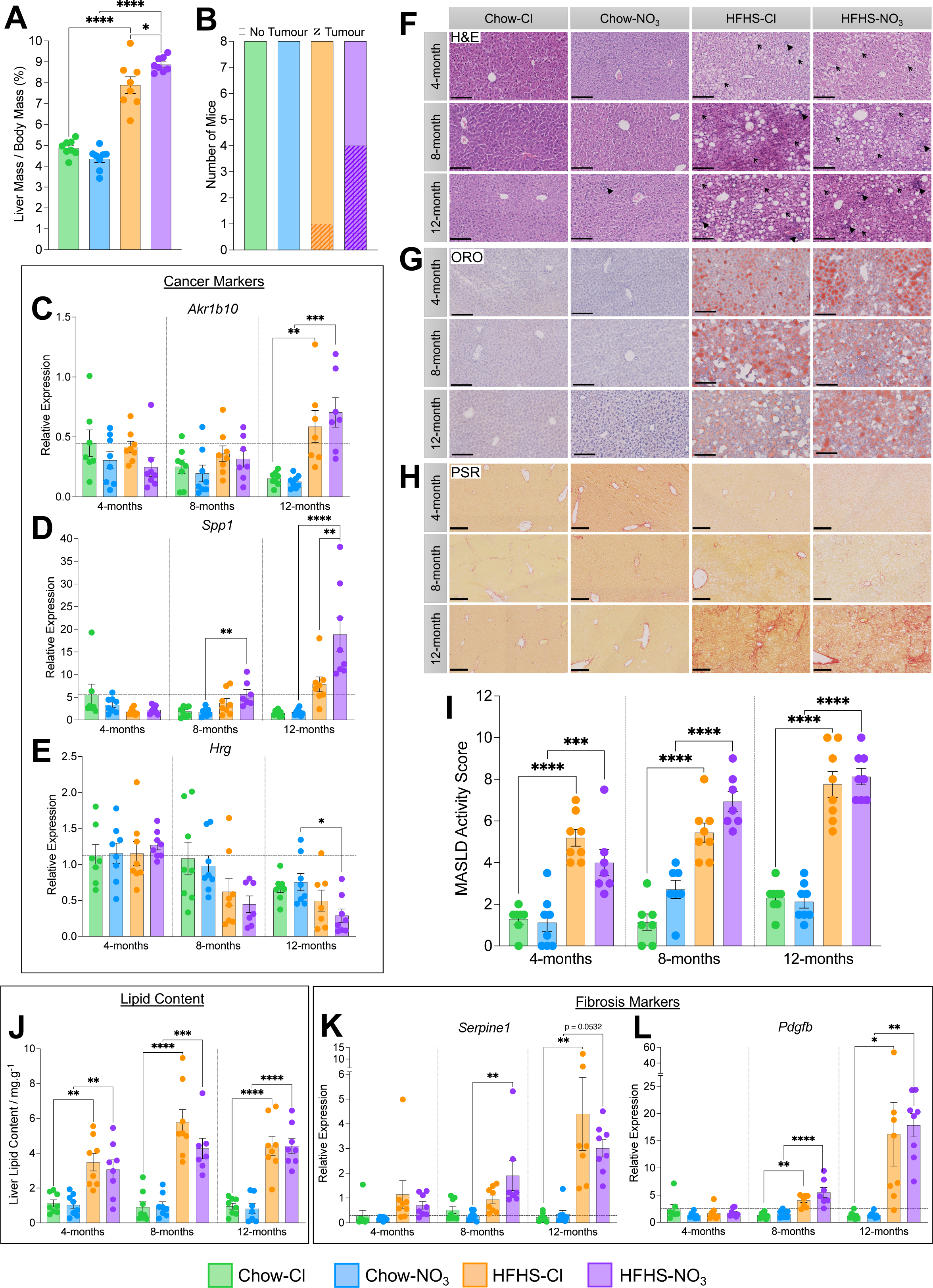
Inorganic nitrate supplementation accelerates the severity of metabolic dysfunction associated steatotic liver disease in mice. (A) Liver mass, normalised to total body mass, in 12-month-old mice. (B) Prevalence of tumors, visible with the naked eye, in livers from 12-month-old-mice. (C) Relative hepatic expression of *Akr1b10*, a marker known to positively associate with hepatocellular carcinoma (HCC) progression, determined by RT-qPCR. Horizontal line indicates relative expression in 4-month-chow/chloride mice. (D) Relative hepatic expression of *Spp1*, a marker known to positively associate with HCC progression, determined by RT-qPCR. Horizontal line indicates relative expression in 4-month-chow/chloride mice. (E) Relative hepatic expression of *Hrg*, a marker known to negatively associate with HCC progression, determined by RT-qPCR. Horizontal line indicates relative expression in 4-month-chow/chloride mice. (F) Representative liver sections stained with hematoxylin and eosin (H&E) at 20x magnification. Scale bars represent 150 mm. Open arrows highlight examples of hepatocyte ballooning; closed arrowheads highlight examples of immune invasion foci. (G) Representative liver sections stained with oil red O (ORO) at 20x magnification. Scale bars represent 150 mm. Neutral lipids and triglycerides are stained red. (H) Representative liver sections stained with picrosirius red (PSR) at 10x magnification. Scale bars represent 250 mm. Collagen fibers are stained red, identifying regions of fibrosis. (I) MASLD activity score (MAS) determined from blinded histological scoring (Suppl. Table 2). (J) Hepatic lipid content determined by modified Folch extraction of lipids. (K) Relative hepatic expression of *Serpine1*, a marker of fibrosis, determined by RT-qPCR. Horizontal line indicates relative expression in 4-month-chow/chloride mice. (L) Relative hepatic expression of *Pdgfb*, a marker of fibrosis, determined by RT-qPCR. Horizontal line indicates relative expression in 4-month-chow/chloride mice. Data represent mean ± SEM; N=7-8 / group. *p<0.05, **p<0.01, ***p<0.001, ****p<0.0001; two-way ANOVA with Tukey’s *post hoc* HSD test for multiple comparisons.

As hepatocellular carcinoma (HCC) is a known consequence of MASLD, we investigated transcriptional markers of HCC (80, 81) in livers across all cohorts (**Fig. 5C-E**). *Akr1b10*, a marker positively associated with HCC progression, was elevated in 12-month-old HFHS-fed mice, compared with chow-fed controls, irrespective of nitrate supplementation (4.7-fold, p<0.01; **Fig. 5C**). Moreover, HFHS-NO_3_-fed mice showed specific elevation of the HCC marker *Spp1*, which was 3.2-fold higher than chow-fed mice at 8-months-of age (p<0.01) and by 12-months, showed expression that was 11.1-fold higher than in chow-fed controls (p<0.0001) and 2.4-fold higher than HFHS-Cl fed mice (p<0.01; **Fig. 5D**). Hepatic *Spp1* expression also showed a progressive increase in HFHS-NO_3_-fed mice, with an 8.6-fold higher (p<0.0001) and 3.3-fold higher (p<0.001) expression at 12-months compared with 4- and 8-months respectively (**Fig. 5D**). Furthermore, *Hrg* (a marker negatively associated with HCC) was lower in livers of HFHS-NO_3_-fed mice from 8-months of age, reaching significance in the 12-month cohort compared with chow-fed controls (61.7%, p<0.05; **Fig. 5E**). *Hrg* expression was also 64.7% lower in 8-month-old mice (p<0.01) and 77.2% lower in 12-month-old mice (p<0.0001) fed a HFHS-NO_3_ diet, compared with the same cohort at 4-months-old (**Fig. 5E**).

We further undertook blind histological scoring of MASLD activity (MAS) in livers from mice of all cohorts (**Fig. 5F-I**). Representative histological images are shown in **Fig. 5F-H**. Mice fed a HFHS diet had livers with MAS that were, on average, 4.46 points higher than those maintained on chow (p<0.001), independent of nitrate supplementation (**Fig. 5I**).

As the steatotic component of MAS was graded 2.1 points higher in HFHS-fed mice compared with chow-fed controls at every time point (p<0.0001, **Suppl. Fig. 4A**), hepatic lipid content was quantified to more accurately examine differences in hepatic lipid load across the experimental cohorts (**Fig. 5J**). Whilst HFHS-fed mice showed clear steatosis with 4.4-fold more fat per gram liver tissue compared with chow-fed controls (p<0.01, **Fig. 5J**), inorganic nitrate supplementation did not alter the steatotic component of MASLD.

In contrast, expression of the fibrotic markers *Serpine1* and *Pdgfb* indicated that fibrosis may be elevated at an earlier stage in HFHS-NO_3_-fed mice compared with HFHS-Cl mice (**Fig. 5K**, **L**). Whilst *Serpine1* expression was elevated 20.0-fold in livers from HFHS-Cl-fed mice at 12-months compared with chow-fed controls, there was no difference in expression of *Serpine1* in HFHS-Cl and chow-Cl livers from younger mice (**Fig. 5K**). In HFHS-fed mice supplemented with NO_3_^-^, however, *Serpine1* showed a 6.7-fold higher expression at 8-months (p<0.05) and 8.5-fold higher expression at 12-months (p = 0.0532) compared with chow-fed controls (**Fig. 5K**). Likewise, *Pdgfb* expression was 3.1-fold higher at 8-months (p<0.01) and 13.3-fold higher (p<0.05) at 12-months in the livers of HFHS-Cl mice compared with those of chow-Cl mice, whilst in HFHS-NO_3_ supplemented mice *Pdgfb* showed a 3.6-fold higher expression at 8-months (p<0.0001) and 13.2-fold higher expression at 12-months (p<0.01) compared with chow-fed controls (**Fig. 5L**).

Taken together, this data suggests that nitrate supplementation accelerates MASLD progression in HFHS-fed mice, including greater incidence of hepatocellular carcinoma. Whilst nitrate did not alter steatosis, fibrosis was initiated earlier in HFHS-NO_3_ livers than those of HFHS-Cl mice.

### Hepatic lipid metabolism and mitochondrial function

Finally, given the changes in steatosis in HFHS-fed mice, we sought to investigate whether the lipidome and, subsequently, fat metabolism, were altered in mice fed a HFHS diet or inorganic nitrate. Open-profiling lipidomics identified over 900 lipid species present in the livers of mice across all experimental conditions and time points. Principal component analysis (PCA) revealed that the hepatic lipidomes of HFHS-fed mice were distinct from those of chow-fed mice (R^2^X_(cum)_ = 88.3%, Q^2^_(cum)_ = 82.2**%**), irrespective of nitrate (**Suppl. Fig. 4B**). Plotting all lipids by species indicated clear differences due to diet in the phosphatidylserine (PS) and TAG complement of the lipidome (**Suppl. Fig. 4C**) with more subtle changes due to age and nitrate supplementation also appearing in these classes.

We therefore investigated the changes occurring in the PS and TAG complement in more depth. Partial least square-discriminant analysis (PLS-DA) of the PS (R^2^X_(cum)_ = 95.3%, R^2^Y_(cum)_ = 40.2%, Q^2^ = 24.1%, p<0.05; **Suppl. Fig. 4D**) and TAG (R^2^X_(cum)_ = 84.6%, R^2^Y_(cum)_ = 28.5%, Q^2^ = 26%, p<0.0001; **Suppl. Fig. 4E**) complements, respectively, in chloride-supplemented mice, revealed that these lipid classes alone were sufficient to discriminate between livers of chow and HFHS-fed mice. PS species were depleted across the board in livers of mice fed a HFHS diet, independent of nitrate supplementation (**Fig. 6A**), and, whilst TAGs were generally increased in HFHS-fed mice livers, there was greatest accumulation of species associated with *de novo* lipogenesis (total carbons 44 to 48, [82]; **Fig. 6B**). Interestingly, the top 5 species differentiating HFHS from chow-fed livers (by variable importance parameter (VIP) score) for both PS and TAGs were highly correlated with the score given for steatosis in MAS (all p<0.0001; Pearson’s correlation). The most highly correlated PS (PS[34:2]; R^2^ = 0.7733) and TAG (TG[46:1]; R^2^ = 0.8211) were 99.9% lower (p<0.0001; **Suppl. Fig. 4F**) and 42.3-fold higher (p<0.0001; **Suppl. Fig. 4F**) respectively in livers with histologically classified steatosis (MAS steatosis score 2 or 3) compared with those without steatosis (MAS steatosis score 0 or 1), suggesting these lipidomic differences are associated with MASLD not just increased dietary intake of lipid.

**Figure 6.**
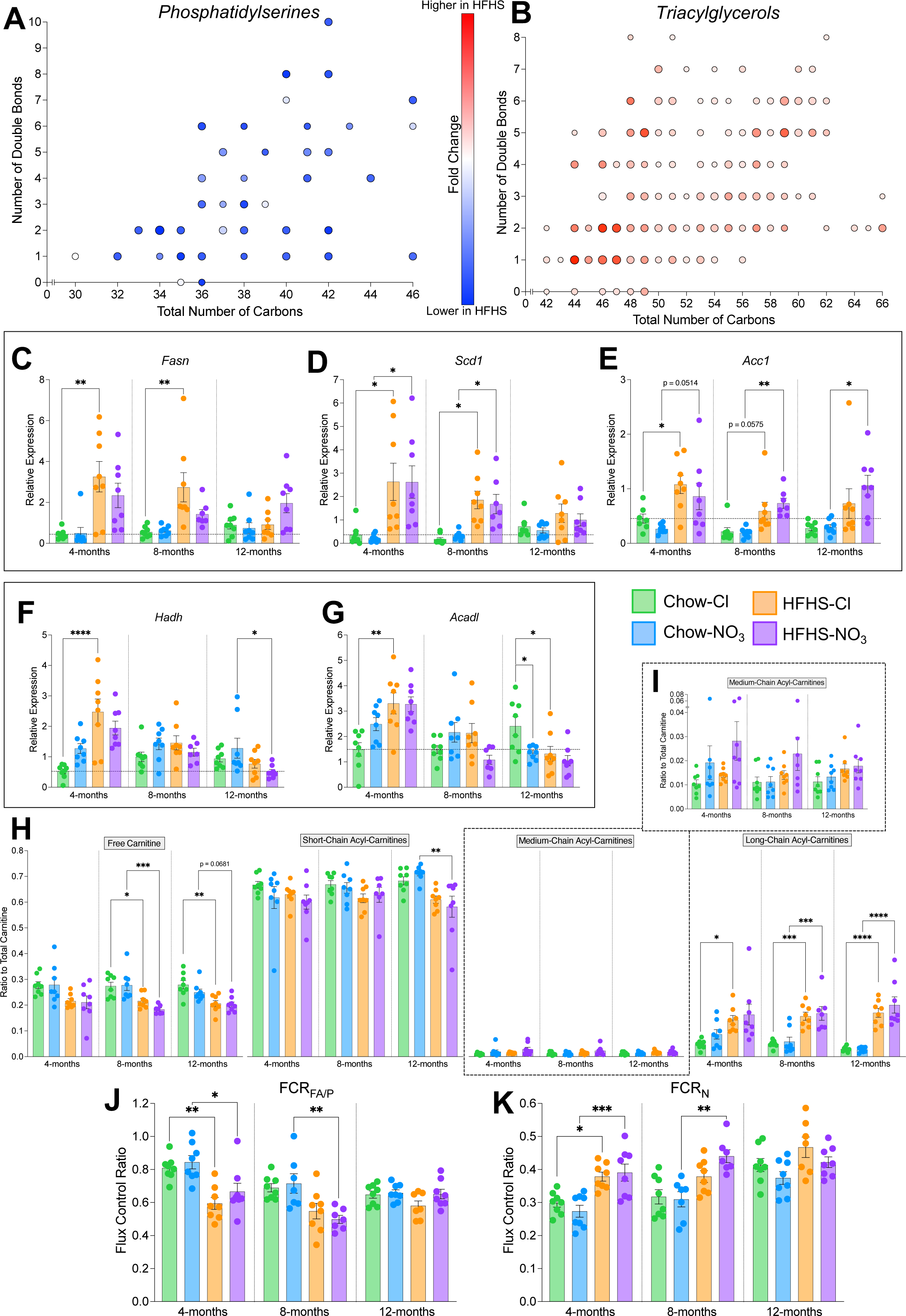
HFHS-feeding is associated with remodeling of the hepatic lipidome and rewiring of lipid metabolism in an age-dependent manner. (A) Hepatic phosphatidylserine (PS) species found to be differentially affected by HFHS-feeding in mice (determined by variable importance parameter (VIP) score from partial least square-discriminant analysis (PLS-DA; **Suppl.** Fig. 4D) greater than 1), plotted by total number of carbons and double bonds in the acyl chains. (B) Hepatic phosphatidylserine (PS) species found to be differentially affected by HFHS-feeding in mice (VIP score from PLS-DA (**Suppl.** Fig. 4E) greater than 1), plotted by total number of carbons and double bonds in the acyl chains. For (A) and (B), size of the point represents the VIP score for the PLS-DA model and color represents relative fold change in livers from HFHS-fed mice compared to chow-fed mice. (C) Relative hepatic expression of *Fasn*, determined by RT-qPCR. Horizontal line indicates relative expression in 4-month-chow/chloride mice. (D) Relative hepatic expression of *Scd1*, determined by RT-qPCR. Horizontal line indicates relative expression in 4-month-chow/chloride mice. (E) Relative hepatic expression of *Acc1*, determined by RT-qPCR. Horizontal line indicates relative expression in 4-month-chow/chloride mice. (F) Relative hepatic expression of *Hadh*, determined by RT-qPCR. Horizontal line indicates relative expression in 4-month-chow/chloride mice. (G) Relative hepatic expression of *Acadl*, determined by RT-qPCR. Horizontal line indicates relative expression in 4-month-chow/chloride mice. (H) Relative levels of free (no associated acyl chain), short-chain (C2-C5), medium-chain (C6-C12) and long-chain (≥C13) acyl-carnitines to the total carnitine pool as measured by liquid chromatography-mass spectrometry (LC-MS) from liver tissue. (I) Relative levels of medium-chain acyl-carnitines to the total carnitine pool (from (H)) shown as an expanded inset. (J) Capacity for OXPHOS supported by octanoyl-carnitine and malate relative to pyruvate and malate, in liver homogenate. (K) Contribution of the N-pathway *via* Complex I to maximal OXPHOS in liver homogenate. For (C)-(K) data represent mean ± SEM; N=7-8 / group. *p<0.05, **p<0.01, ***p<0.001, ****p<0.0001; two-way ANOVA with Tukey’s *post hoc* HSD test for multiple comparisons.

We next considered the expression of genes involved in hepatic lipid catabolism and anabolism. Lipogenesis genes were found to be generally upregulated in HFHS-fed mice (**Fig. 6C-E**). Fatty acid synthase (*Fasn*) showed a 7.4-fold higher expression at 4-months and 5.0-fold higher expression at 8-months in HFHS-Cl mice compared with chow-fed controls (p<0.01; **Fig. 6C**) with no impact of inorganic nitrate supplementation. Stearoyl-coA desaturase 1 (*Scd1*) was expressed 8.1-fold higher in livers from 4- and 8-month-old HFHS-fed mice regardless of nitrate-supplementation (p<0.05; **Fig. 6D**). Similarly, acetyl-coA carboxylase (*Acc1*) was expressed 2.9-fold higher in 4-and 8-month-old HFHS-fed mice (p£0.05; **Fig. 6E**), but remained elevated in 12-month-old HFHS-NO_3_ mice, where expression was 3.6-fold higher than in chow-fed controls (p<0.05; **Fig. 6E**). Interestingly, the genes encoding the β-oxidation enzymes 3-hydroxyacyl-CoA dehydrogenase (*Hadh*) and long-chain acyl-coA dehydrogenase (*Acadl*) showed an age-dependent expression profile (**Fig. 6F**, **G**). In 4-month-old HFHS-Cl mice, *Hadh* expression was 4.7-fold higher (p<0.0001; **Fig. 6F**) and *Acadl* expression was 2.2-fold higher (p<0.01; **Fig. 6G**) compared with chow-fed controls. However, there was no difference due to diet nor nitrate-supplementation in 8-month-old mice, and by 12-months-old, *Hadh* expression was 60.3% lower in HFHS-NO_3_ mice (p<0.05; **Fig. 6F**) and *Acadl* expression was 45.4% lower in HFHS-Cl mice (p<0.05, **Fig. 6G**) compared with chow-fed controls. Chow-NO_3_-fed mice also showed 44.0% lower *Acadl* expression at 12-months compared with chow-Cl-fed mice (p<0.05; **Fig. 6G**).

We next investigated the acyl-carnitine profile of the liver. The total acyl-carnitine pool was 40.7% lower in HFHS-fed mice compared with chow-fed mice (**Suppl. Fig. 5A**). However, as in the heart, there were alterations to the composition of the hepatic acyl-carnitine pool independent of nitrate supplementation (**Fig. 6H**, **I**; **Suppl. Fig. 5B**). At all time-points, relative levels of long-chain (³C13) acyl-carnitines were 3.4-fold higher in HFHS-fed mice (p<0.05), whilst 8- and 12-month-old HFHS-fed mice showed 22.8% lower contribution of free carnitine to the total carnitine pool (**Fig. 6H**). Surprisingly, there was no concurrent suppression in the relative level of short-chain (C2-C5) acyl-carnitines with the accumulation in long-chain acyl-carnitines, except in 12-month-old HFHS-NO_3_ mice, where the relative contribution of short-chain acyl-carnitines to the total carnitine pool was 18.2% lower than chow-fed controls (p<0.01; **Fig. 6H**). There were no differences in relative levels of medium-chain acyl-carnitines (**Fig. 6H**, **I**).

Finally, we assessed whether changes measured in the liver during MASLD progression were associated with alterations in mitochondrial respiratory function. Generally, mice fed a HFHS-diet tended to have a lower mass-corrected respiratory capacity in all states at all ages, independent of nitrate supplementation. This was especially evident with β-oxidation supported respiration (OctM*_L_* and OctM*_P_*) and following stimulation of complex II with succinate (GMS*_P_*, GMS*_E_* and S*_E_*; **Suppl. Fig. 5C**). In-line with this, the capacity for fatty acid oxidation relative to pyruvate oxidation (FCR_FA/P_) was 23.7% lower in the livers of 4-month-old HFHS-fed mice (p<0.05) and 30.3% lower in the livers of 8-month-old HFHS-NO_3_ mice (p<0.01), compared with chow-fed controls (**Fig. 6J**). However, the contribution of the N-pathway to maximal OXPHOS (FCR_N_) was 34.6% higher in 4-month-old HFHS-fed mice regardless of nitrate supplementation (p<0.05) and 42.1% higher in 4-month-old HFHS-NO_3_ mice (p<0.01) compared with chow-fed controls (**Fig. 6K**).

Taken together, these results highlight that hepatic lipid metabolism is impacted by HFHS-feeding, with metabolic changes occurring early and persisting throughout life in mice fed a HFHS-diet.

## Discussion

Inorganic nitrate has been suggested as a possible treatment for obesity-related metabolic disease owing to the relative ease of modifying intake through the diet (83) and favorable effects upon acute administration including increased bioavailability of NO (20), improvement of mitochondrial efficiency (66) and enhanced mitochondrial β-oxidation (35). Collectively, these have been proposed to be beneficial in treating metabolic diseases by ameliorating any effects of mitochondrial dysfunction and providing a mechanism for dissipating excess FFA. However, here we show that dietary supplementation with a moderate dose of inorganic nitrate supplementation (achievable in humans *via* a modest increase in leafy green vegetable consumption [84]) was not effective as a therapeutic intervention in mice with diet-induced obesity, and was instead associated with adverse effects in HFHS-fed mice (**Fig. 7**). Most notably, dietary supplementation with inorganic nitrate resulted in an elevated plasma LDL cholesterol, increased cardiac fibrosis and accelerated MASLD severity in HFHS-fed mice, including a significant, detectable tumor burden in 12-month-old HFHS-NO_3_ fed mice.

**Figure 7.**
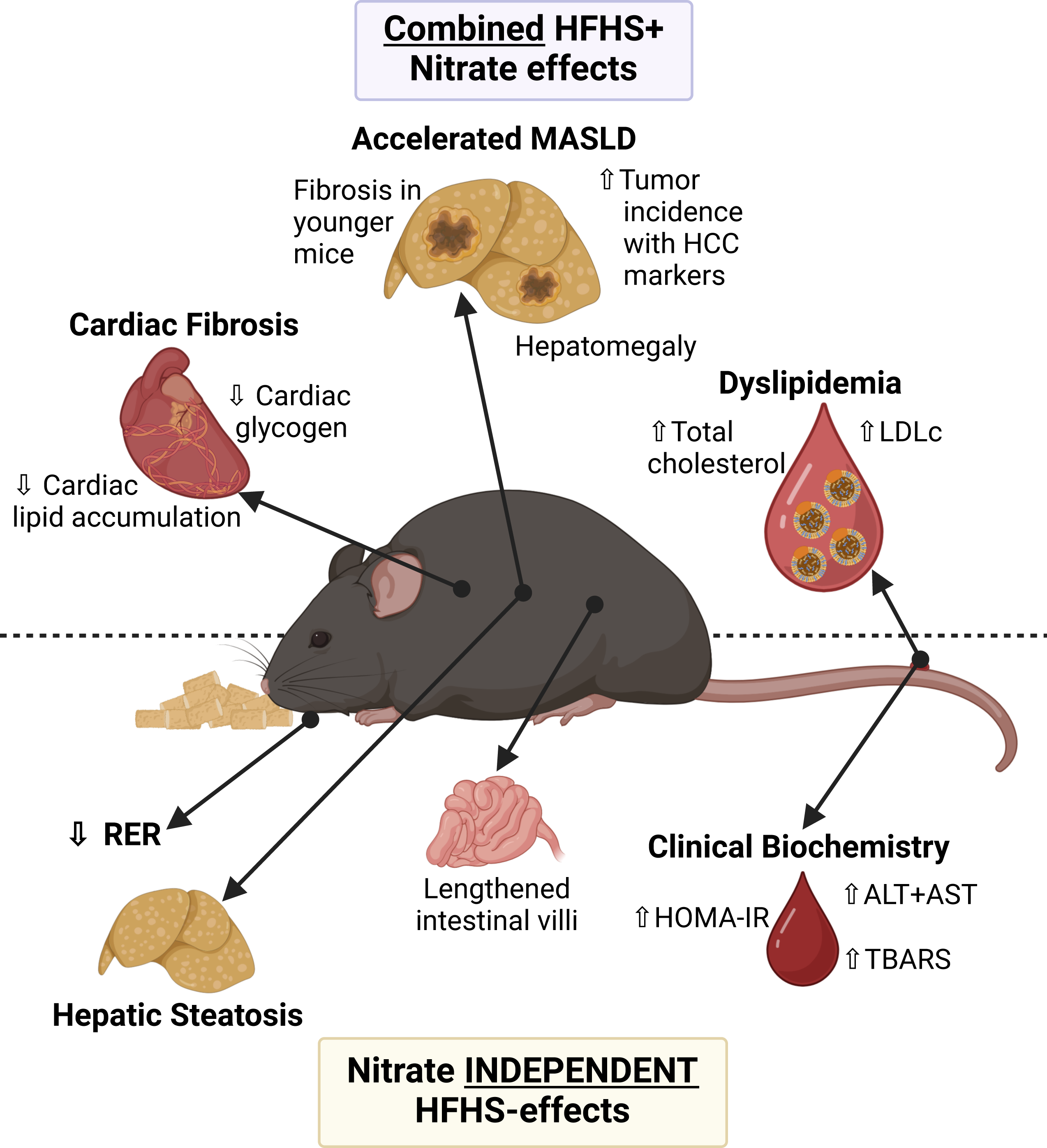
Summary of diet-induced obesity effects determined in this study as a result of HFHS-feeding (independent of nitrate supplementation) or as a result of inorganic nitrate supplementation in combination with HFHS-diet. ↑ indicates higher in HFHS (compared with chow) or HFHS-NO_3_ (compared with HFHS-Cl) respectively; ↓ indicates lower in HFHS (compared with chow) or HFHS-NO_3_ (compared with HFHS-Cl) respectively. *ALT – alanine transaminase; AST – aspartate transaminase; HCC – hepatocellular carcinoma; HOMA-IR – homeostatic model assessment of insulin resistance; LDLc – low density lipoprotein cholesterol; MASLD – metabolic-dysfunction associated steatotic liver disease; RER – respiratory exchange ratio; TBARS – thiobarbituric acid reactive substances.* Created with BioRender.com.

A major strength of this study is that we have investigated the metabolic phenotype at multiple timepoints over a sustained timeframe. Many studies investigating metabolic alterations bought about by obesity or diabetes consider a single time point in comparison with a healthy control. However, type-II diabetes is a progressive disease that exists on a spectrum from mild insulin resistance to overt glucose intolerance with b-cell impairment and it is therefore important to consider when and where adverse metabolic remodeling occurs as the disease progresses. This temporal analysis has also allowed us to consider some of the longer-term effects associated with nitrate-supplementation, which are not apparent over a shorter time frame. Furthermore, we have utilized a range of gold-standard techniques allowing an in-depth dissection of the progressive obesity phenotype alongside changes in mitochondrial function, tissue metabolism and morphology.

Our findings add clarity to the time course of metabolic derailment occurring over the development of obesity and related comorbidities. This is evident, for instance, in the liver where our in-depth analysis of the MASLD phenotype, accompanied with whole lipidome analysis and mitochondrial functional measurements has allowed a comprehensive assessment of the metabolic changes occurring from a healthy tissue all the way to severe disease. For instance, whilst PS depletion has previously been shown to occur in MASH patients (85), our data highlights that PS depletion occurs from the very earliest time points of metabolic derailment, prior to progression to MASH. Furthermore, combined suppression of PSS1 and PSS2 (enzymes involved in the synthesis of PS) lowered hepatic PS in mice and concurrently increased hepatic TAG (86), in agreement with the changes reported here. However, we additionally show that these changes are accompanied by alterations in the balance between hepatic lipid catabolism and anabolism, alongside changes in mitochondrial respiratory function.

Our finding of detrimental consequences of inorganic nitrate in HFHS-diet fed mice (**Fig. 7**) can be considered alongside previously published benefits of nitrate supplementation in healthy rodents. For instance, up to 17-months of inorganic nitrate supplementation at an equivalent dose was shown to have no adverse effects in male C57Bl/6 mice, and improved fasting insulin in these mice (42), however, these mice were maintained on a standard laboratory chow (with depleted nitrate, [42]). The adverse effects observed in the HFHS-NO_3_ mice here are likely to result from an interaction between the obesogenic diet and the nitrate itself; indeed, we also observed no adverse effects in nitrate supplemented mice fed a standard laboratory chow diet. Of note, however, nitrate-supplementation did not result in any improvement in fasting insulin in chow-fed mice in our study, contrary to that previously observed (42).

Furthermore, inorganic nitrate has been shown to increase mitochondrial β-oxidation capacity in rat skeletal muscle (35, 87), a finding which gave rise to some of the hypothesized benefits of inorganic nitrate supplementation in obesity-related comorbidities. However, this effect was not observed in NO_3_-supplemented mice here, regardless of whether they received standard chow or HFHS diet, or whether soleus or gastrocnemius muscle was analyzed. This apparent discrepancy is likely due to the lower baseline nitrate levels of rats in comparison to mice (88) resulting in reduced elevations in plasma nitrate levels achievable in mice, as used in this study, through dietary supplementation with NaNO_3_. Whilst plasma levels of nitrate in rats are more similar to those in humans (88, 89), one of the significant advantages of this work was that by using mice rather than rats, a long-term study could be carried out (90), allowing us to observe the metabolic alterations over the entire course of obesity progression.

Additionally, our finding of increased cardiac fibrosis in HFHS-NO_3_ mice contrasts with previous studies which have shown a nitrate driven amelioration of fibrosis in the kidney and heart as a result of obesity or hypertension in rodent models (91–94). This apparent discrepancy may be attributable to the significantly longer timescale investigated here, compared with a maximum of 10-weeks in previous studies. Alternatively, an endothelial-to-mesenchymal transition due to inflammation or a perturbation in redox homeostasis, along with the known reduced effectiveness of the Nrf2 system (and thus fibrosis resolution) with aging (95, 96) may account for the increased cardiac fibrosis seen here. This elevation in cardiac fibrosis was particularly surprising to find in HFHS-NO_3_ mice, given the anti-inflammatory effects purported to nitro-fatty acids (nitro-FAs); endogenously produced electrophilic metabolites resulting from the non-enzymatic reaction between unsaturated fatty acids and NO/NO_3_^-^ oxidation products (97, 98). Whilst the HFHS diet given to these mice was predominantly high in saturated fats, the unsaturated fat content was thrice that of the standard laboratory chow (6.6% vs. 2.2% *w/w*, according to manufacturer analyses), which would be suspected to result in increased nitro-FAs concomitantly. However, as the anti-inflammatory effects of nitro-FAs occur, at least partially, through activation of the Nrf2 system (98, 99), perhaps the age-related decline in effectiveness also accounts for the surprising increase in fibrosis. If not, there may perhaps be an alteration of the balance of nitro-FA formation and the relationship with Nrf2 mediated processes that leads to an unfavorable shift in the wrong direction in HFHS-NO_3_ fed mice.

An unexpected finding in this study was the increased incidence of liver tumors in 12-month HFHS-NO_3_ fed mice with gene expression analysis suggesting that concurrent HFHS-NO_3_ feeding accelerated progression from MASLD to HCC. The pathogenesis of MASLD-associated HCC is incompletely understood, as, despite MASLD being the most common underlying risk factor for HCC (100), only a minority of MASLD patients go on to develop HCC (101). The current hypothesized factors underlying the increased HCC risk in MASLD include compensatory hyperproliferation to counter hepatocyte cell death (as a result of lipotoxicity or DNA damage from oxidative damage), activation of hepatic stellate cells resulting in increased fibrosis, and elevated inflammatory factors (101). Whilst inorganic nitrate was administered to mice at a low dose, it was expected to increase NO bioavailability, which, whilst not considered to be damaging to DNA alone, can produce reactive nitrogen species (RNS) upon reacting with reactive oxygen species (ROS) that can damage DNA through both base mutations and strand breaks (102). MASLD has been associated with increased ROS production as a result of fatty acid overload, which may reduce mitochondrial coupling and increase inner mitochondrial membrane electron leak (103). The combined increase of ROS (as a result of the HFHS diet) and NO (as a result of dietary nitrate) may therefore be a key event precipitating the increased incidence of apparent HCC with HFHS-NO_3_ feeding. Although the reduction of nitrate to nitrite is likely slow and quantitatively moderate, tissue nitrite may exceed levels in the circulation (88); thus we cannot exclude the possibility that liver tumors may be induced as a result of enhanced nitrosamine formation.

Several studies have highlighted the potential benefits of nitrate supplementation in obesity-related metabolic diseases (28, 31–33, 104). Much of this work stems from the initial observation that dietary inorganic nitrate increased NO bioavailability, offsetting the fall in endogenous NO production, which may be a key pathological event in the development of MetS (28). However, these experiments were carried out on eNOS-deficient mice (28), raising the possibility that nitrate may only be beneficial against the background of reduced endogenous NO production. Inorganic nitrate supplementation may be less effective in cases of MetS or T2DM where endogenous NOS enzymes are functional, as, although there can be impaired *eNOS* activity in MetS and T2DM (105, 106), there is a more subtle endogenous NO-depletion than that seen in *eNOS*-deficient mice. Support for this arises from the observation that 12-week nitrate supplementation did not ameliorate MetS development in high-fat diet-fed mice (107), much like nitrate supplementation did not improve adiposity or HOMA-IR in the HFHS-fed mice in this study.

Despite these negative findings, nitrate may yet hold benefit as a therapeutic agent in certain individuals with obesity-related metabolic disease. *eNOS* polymorphisms have been associated with T2DM and MetS (14, 15), highlighting a group of patients in which nitrate supplementation may prove to be a beneficial treatment strategy. Furthermore, a higher intake of leafy green vegetables (the largest dietary source of inorganic nitrate [(108)]) is associated with a 14% reduced risk of T2DM development (103). Whilst this may be a result of general dietary habits (i.e., individuals who consume more leafy vegetables may consume less processed foods) it highlights the possibility of a preventative role of nitrate in T2DM development as opposed to using it as a therapeutic agent for the treatment of the condition.

Nevertheless, our work highlights that nitrate supplementation is largely ineffective in protecting against the significant consequences of long-term HFHS-feeding. Further, we have uncovered that negative consequences of nitrate can emerge against the background of an obesogenic diet. Thus, nitrate supplementation is unlikely to be an effective approach to improve metabolic health in obesity, without wider alterations to the diet. Given the recent change in public perception of nitrate as being beneficial rather than detrimental, there is an urgent need to reproduce our findings in other laboratories and animal models and to investigate the mechanisms of enhanced cardiac fibrosis and hepatic malignancy. This is especially important since epidemiological evidence is emerging that the effects of enhanced nitrate intake on cardiovascular health may be source-specific (109).

## Data Availability

The datasets supporting the results presented in this article will be made freely available *via* the Cambridge University Repository upon acceptance of publication. Raw metabolomics data will be deposited to the EMBL-EBI MetaboLights database upon acceptance of publication.

## Supplemental Material

Supplemental methods, five supplemental figures and two supplemental tables accompany the manuscript.

## Acknowledgments

For the purpose of open access, the author has applied a Creative Commons Attribution (CC BY) licence to any Author Accepted Manuscript version arising from this submission.

We thank Sarah Royle, Matthew Rodgers and Mithylan Ganeshwaran for their help on the project and Benjamin Stockell for statistical advice. We also thank Hayley Forrest, Alison Robertson and Ben Jaggs for support with animal work and Joe Lewis for support with the metabolic cage set-up. We thank Kat Millen and Sarah Bray for use of thermocycler equipment throughout the study. We thank Hannah Mannering and the Disease Model Core (Wellcome-MRC Institute of Metabolic Sciences) for their technical assistance in DEXA scans, James Warner and the Histopathology Core (MRC Metabolic Diseases Unit [MC_UU_00014/5]) for processing and embedding liver samples and Peter Barker, Keith Burling and the Cambridge Biochemical Assay Laboratory (Cambridge University Hospitals NHS Foundation Trust) for clinical chemistry analyses.

We acknowledge the following grants for support and funding the work presented in this manuscript: 4-year PhD studentships from the British Heart Foundation, Grant Numbers: FS/17/61/33473 (to APS); FS/4yPhD/F/21/34156 (to RB); FS/18/56/34177 (to DKK); FS/4yPhD/F/20/34124 (to BDT). Rank Prize Fund Nutrition Covid Response Grant (to APS).

4-year PhD studentship from the Wellcome Trust, Grant Number: 220033/Z/19/Z (to LMWH).

Biotechnology and Biological Sciences Research Council Doctoral Training Program, Grant Number: BB/M011194/1 (to PMD).

Medical Research Council Lipid Profiling and Signalling & Lipid Dynamics and Regulation, Grants Numbers: MC UP A90 1006 and MC PC 13030 (to GM, MV and JLG)

Research Councils UK, Grant Number: EP/E500552/1 (to AJM)

Evelyn Trust, Grant Number: 16/33 (to AJM).

## Present addresses

Present address of A.P.S: Medical Research Council Mitochondrial Biology Unit, University of Cambridge, Cambridge, CB2 0XY, UK.

Present address of R.B: Department of Pharmacology, University of Cambridge, Cambridge, CB2 1PD, UK.

Present address of G.M and M.V: Roger Williams Institute of Hepatology, Foundation for Liver Research, London, SE5 9NT, UK.

Present address of M.V: Department of Interdisciplinary Medicine, Clinica Medica “C. Frugoni”, Aldo

Moro University of Bari, 70124 Bari, Italy.

## Disclosures

The authors declare no conflicts of interest.

## Author contributions

APS, JLG and AJM conceived and designed research with input and advice from MV and MF. APS, LMWH, FNK, RB, GM, DKK, MM, KAO, MCH, PMD and BDT performed experiments. APS analyzed data, interpreted results of experiments, prepared figures and drafted manuscript. MF, JLG and AJM edited and revised the manuscript. All authors approved the final version of the manuscript.

## Notes

### Competing Interest Statement

The authors have declared no competing interest.

## References

1. The GBD 2015 Obesity Collaborators. Health Effects of Overweight and Obesity in 195 Countries over 25 Years. N Engl J Med 377: 13–27, 2017. doi: 10.1056/NEJMoa1614362.

2. The Global BMI Mortality Collaboration. Body-mass index and all-cause mortality: individual-participant-data meta-analysis of 239 prospective studies in four continents. Lancet 388: 776–786, 2016. doi: 10.1016/S0140-6736(16)30175-1.

3. Engin A. The Definition and Prevalence of Obesity and Metabolic Syndrome. In: Obesity and Lipotoxicity, edited by Engin AB, Engin A. Springer International Publishing, p. 1–17.

4. Wang YC, McPherson K, Marsh T, Gortmaker SL, Brown M. Health and economic burden of the projected obesity trends in the USA and the UK. Lancet 378: 815–825, 2011. doi: 10.1016/S0140-6736(11)60814-3.

5. Kahn SE, Hull RL, Utzschneider KM. Mechanisms linking obesity to insulin resistance and type 2 diabetes. Nature 444: 840–846, 2006. doi: 10.1038/nature05482.

6. Kim MS, Kim WJ, Khera AV, Kim JY, Yon DK, Lee SW, Shin JI, Won H-H. Association between adiposity and cardiovascular outcomes: an umbrella review and meta-analysis of observational and Mendelian randomization studies. Eur Heart J 42: 3388–3403, 2021. doi: 10.1093/eurheartj/ehab454.

7. Milić S, Lulić D, Štimac D. Non-alcoholic fatty liver disease and obesity: Biochemical, metabolic and clinical presentations. World J Gastroenterol 20: 9330–9337, 2014. doi: 10.3748/wjg.v20.i28.9330.

8. Weidmann P, Boehlen LM, de Courten M. Pathogenesis and treatment of hypertension associated with diabetes mellitus. Am Heart J 125: 1498–1513, 1993. doi: 10.1016/0002-8703(93)90447-H.

9. Huang PL, Huang Z, Mashimo H, Bloch KD, Moskowitz MA, Bevan JA, Fishman MC. Hypertension in mice lacking the gene for endothelial nitric oxide synthase. Nature 377: 239–242, 1995. doi: 10.1038/377239a0.

10. Duplain H, Burcelin R, Sartori C, Cook S, Egli M, Lepori M, Vollenweider P, Pedrazzini T, Nicod P, Thorens B, Scherrer U. Insulin Resistance, Hyperlipidemia, and Hypertension in Mice Lacking Endothelial Nitric Oxide Synthase. Circulation 104: 342–345, 2001. doi: 10.1161/01.CIR.104.3.342.

11. Cook S, Hugli O, Egli M, Vollenweider P, Burcelin R, Nicod P, Thorens B, Scherrer U. Clustering of cardiovascular risk factors mimicking the human metabolic syndrome X in *eNOS* null mice. Swiss Med Wkly 133: 360–363, 2003. doi: 10.4414/smw.2003.10239.

12. Nisoli E, Clementi E, Paolucci C, Cozzi V, Tonello C, Sciorati C, Bracale R, Valerio A, Francolini M, Moncada S, Carruba MO. Mitochondrial Biogenesis in Mammals: The Role of Endogenous Nitric Oxide. Science 299: 896–899, 2003. doi: 10.1126/science.1079368.

13. Le Gouill E, Jimenez M, Binnert C, Jayet P-Y, Thalmann S, Nicod P, Scherrer U, Vollenweider P. Endothelial Nitric Oxide Synthase (*eNOS*) Knockout Mice Have Defective Mitochondrial β-Oxidation. Diabetes 56: 2690–2696, 2007. doi: 10.2337/db06-1228.

14. Monti LD, Barlassina C, Citterio L, Galluccio E, Berzuini C, Setola E, Valsecchi G, Lucotti P, Pozza G, Bernardinelli L, Casari G, Piatti P. Endothelial Nitric Oxide Synthase Polymorphisms Are Associated With Type 2 Diabetes and the Insulin Resistance Syndrome. Diabetes 52: 1270–1275, 2003. doi: 10.2337/diabetes.52.5.1270.

15. Fernandez ML, Ruiz R, Gonzalez MA, Ramirez-Lorca R, Couto C, Ramos A, Gutierrez-Tous R, Rivera JM, Ruiz A, Real LM, Grilo A. Association of NOS3 gene with metabolic syndrome in hypertensive patients. Thromb Haemost 92: 413–418, 2004. doi: 10.1160/TH04-02-0103.

16. Avogaro A, Toffolo G, Kiwanuka E, de Kreutzenberg SV, Tessari P, Cobelli C. L-Arginine-Nitric Oxide Kinetics in Normal and Type 2 Diabetic Subjects: A Stable-Labelled ^15^N Arginine Approach. Diabetes 52: 795–802, 2003. doi: 10.2337/diabetes.52.3.795.

17. Lundberg JO, Govoni M. Inorganic nitrate is a possible source for systemic generation of nitric oxide. Free Radic Biol Med 37: 395–400, 2004. doi: 10.1016/j.freeradbiomed.2004.04.027.

18. Duncan C, Dougall H, Johnston P, Green S, Brogan R, Leifert C, Smith L, Golden M, Benjamin N. Chemical generation of nitric oxide in the mouth from the enterosalivary circulation of dietary nitrate. Nat Med 1: 546–551, 1995. doi: 10.1038/nm0695-546.

19. Lundberg JO, Weitzberg E, Lundberg JM, Alving K. Intragastric nitric oxide production in humans: measurements in expelled air. Gut 35: 1543–1546, 1994. doi: 10.1136/gut.35.11.1543.

20. Lundberg JO, Weitzberg E, Gladwin MT. The nitrate–nitrite–nitric oxide pathway in physiology and therapeutics. Nat Rev Drug Discov 7: 156–167, 2008. doi: 10.1038/nrd2466.

21. Zhang Z, Naughton DP, Blake DR, Benjamin N, Stevens CR, Winyard PG, Symons MCR, Harrison R. Human xanthine oxidase converts nitrite ions into nitric oxide (NO). Biochem Soc Trans 25: 524S, 1997. doi: 10.1042/bst025524s.

22. Cosby K, Partovi KS, Crawford JH, Patel RP, Reiter CD, Martyr S, Yang BK, Waclawiw MA, Zalos G, Xu X, Huang KT, Shields H, Kim-Shapiro DB, Schechter AN, Cannon RO, Gladwin MT. Nitrite reduction to nitric oxide by deoxyhemoglobin vasodilates the human circulation. Nat Med 9: 1498–1505, 2003. doi: 10.1038/nm954.

23. Shiva S, Huang Z, MacArthur PH, Ringwood LA, Gladwin MT. Myoglobin Is a Nitrite Reductase That Generates NO and Regulates Mitochondrial Respiration. Blood 108: 1561–1561, 2006. doi: 10.1182/blood.V108.11.1561.1561.

24. Moncada S, Higgs A. The L-Arginine-Nitric Oxide Pathway. N Engl J Med 329: 2002–2012, 1993. doi: 10.1056/NEJM199312303292706.

25. Sindelar JJ, Milkowski AL. Human safety controversies surrounding nitrate and nitrite in the diet. Nitric Oxide 26: 259–266, 2012. doi: 10.1016/j.niox.2012.03.011.

26. Zweier JL, Wang P, Samouilov A, Kuppusamy P. Enzyme-independent formation of nitric oxide in biological tissues. Nat Med 1: 804–809, 1995. doi: 10.1038/nm0895-804.

27. Butler AR, Feelisch M. Therapeutic Uses of Inorganic Nitrite and Nitrate. Circulation 117: 2151– 2159, 2008. doi: 10.1161/CIRCULATIONAHA.107.753814.

28. Carlström M, Larsen FJ, Nyström T, Hezel M, Borniquel S, Weitzberg E, Lundberg JO. Dietary inorganic nitrate reverses features of metabolic syndrome in endothelial nitric oxide synthase-deficient mice. Proc Natl Acad Sci U S A 107: 17716–17720, 2010. doi: 10.1073/pnas.1008872107.

29. Jiang H, Torregrossa AC, Potts A, Pierini D, Aranke M, Garg HK, Bryan NS. Dietary nitrite improves insulin signaling through GLUT4 translocation. Free Radic Biol Med 67: 51–57, 2014. doi: 10.1016/j.freeradbiomed.2013.10.809.

30. Singamsetty S, Watanabe Y, Guo L, Corey C, Wang Y, Tejero J, McVerry BJ, Gladwin MT, Shiva S, O’Donnell CP. Inorganic nitrite improves components of the metabolic syndrome independent of weight change in a murine model of obesity and insulin resistance. J Physiol 593: 3135–3145, 2015. doi: 10.1113/JP270386.

31. Essawy SS, Abdel-Sater KA, Elbaz AA. Comparing the effects of inorganic nitrate and allopurinol in renovascular complications of metabolic syndrome in rats: role of nitric oxide and uric acid. Arch Med Sci 10: 537–545, 2014. doi: 10.5114/aoms.2013.33222.

32. Gheibi S, Jeddi S, Carlström M, Gholami H, Ghasemi A. Effects of long-term nitrate supplementation on carbohydrate metabolism, lipid profiles, oxidative stress, and inflammation in male obese type 2 diabetic rats. Nitric Oxide 75: 27–41, 2018. doi: 10.1016/j.niox.2018.02.002.

33. Li T, Lu X, Sun Y, Yang X. Effects of spinach nitrate on insulin resistance, endothelial dysfunction markers and inflammation in mice with high-fat and high-fructose consumption. Food Nutr Res 60: 10.3402/fnr.v60.32010, 2016. doi: 10.3402/fnr.v60.32010.

34. Ashmore T, Fernandez BO, Branco-Price C, West JA, Cowburn AS, Heather LC, Griffin JL, Johnson RS, Feelisch M, Murray AJ. Dietary nitrate increases arginine availability and protects mitochondrial complex I and energetics in the hypoxic rat heart. J Physiol 592: 4715–4731, 2014. doi: 10.1113/jphysiol.2014.275263.

35. Ashmore T, Roberts LD, Morash AJ, Kotwica AO, Finnerty J, West JA, Murfitt SA, Fernandez BO, Branco C, Cowburn AS, Clarke K, Johnson RS, Feelisch M, Griffin JL, Murray AJ. Nitrate enhances skeletal muscle fatty acid oxidation via a nitric oxide-cGMP-PPAR-mediated mechanism. BMC Biol 13: 110, 2015. doi: 10.1186/s12915-015-0221-6.

36. Horscroft JA, O’Brien KA, Clark AD, Lindsay RT, Steel AS, Procter NEK, Devaux J, Frenneaux M, Harridge SDR, Murray AJ. Inorganic nitrate, hypoxia, and the regulation of cardiac mitochondrial respiration—probing the role of PPARα. FASEB J 33: 7563–7577, 2019. doi: 10.1096/fj.201900067R.

37. Ivarsson N, Schiffer TA, Hernández A, Lanner JT, Weitzberg E, Lundberg JO, Westerblad H. Dietary nitrate markedly improves voluntary running in mice. Physiol Behav 168: 55–61, 2017. doi: 10.1016/j.physbeh.2016.10.018.

38. Roberts LD, Ashmore T, Kotwica AO, Murfitt SA, Fernandez BO, Feelisch M, Murray AJ, Griffin JL. Inorganic Nitrate Promotes the Browning of White Adipose Tissue Through the Nitrate-Nitrite-Nitric Oxide Pathway. Diabetes 64: 471–484, 2015. doi: 10.2337/db14-0496.

39. Peleli M, Ferreira DMS, Tarnawski L, McCann Haworth S, Xuechen L, Zhuge Z, Newton PT, Massart J, Chagin AS, Olofsson PS, Ruas JL, Weitzberg E, Lundberg JO, Carlström M. Dietary nitrate attenuates high-fat diet-induced obesity via mechanisms involving higher adipocyte respiration and alterations in inflammatory status. Redox Biol 28: 101387, 2020. doi: 10.1016/j.redox.2019.101387.

40. McNally BD, Moran A, Watt NT, Ashmore T, Whitehead A, Murfitt SA, Kearney MT, Cubbon RM, Murray AJ, Griffin JL, Roberts LD. Inorganic Nitrate Promotes Glucose Uptake and Oxidative Catabolism in White Adipose Tissue Through the XOR-Catalyzed Nitric Oxide Pathway. Diabetes 69: 893–901, 2020. doi: 10.2337/db19-0892.

41. Tateya S, Rizzo NO, Handa P, Cheng AM, Morgan-Stevenson V, Daum G, Clowes AW, Morton GJ, Schwartz MW, Kim F. Endothelial NO/cGMP/VASP Signaling Attenuates Kupffer Cell Activation and Hepatic Insulin Resistance Induced by High-Fat Feeding. Diabetes 60: 2792–2801, 2011. doi: 10.2337/db11-0255.

42. Hezel MP, Liu M, Schiffer TA, Larsen FJ, Checa A, Wheelock CE, Carlström M, Lundberg JO, Weitzberg E. Effects of long-term dietary nitrate supplementation in mice. Redox Biol 5: 234– 242, 2015. doi: 10.1016/j.redox.2015.05.004.

43. Weir JB de V. New methods for calculating metabolic rate with special reference to protein metabolism. J Physiol 109: 1–9, 1949. doi: 10.1113/jphysiol.1949.sp004363.

44. Virtue S, Lelliott CJ, Vidal-Puig A. What is the most appropriate covariate in ANCOVA when analysing metabolic rate? Nat Metab 3: 1585–1585, 2021. doi: 10.1038/s42255-021-00505-5.

45. Mina AI, LeClair RA, LeClair KB, Cohen DE, Lantier L, Banks AS. CalR: A Web-Based Analysis Tool for Indirect Calorimetry Experiments. Cell Metab 28: 656–666.e1, 2018. doi: 10.1016/j.cmet.2018.06.019.

46. R Core Team. R: A language and environment for statistical computing. [Online]. R Foundation for Statistical Computing. https://www.R-project.org/.

47. Friedewald WT, Levy RI, Fredrickson DS. Estimation of the concentration of low-density lipoprotein cholesterol in plasma, without use of the preparative ultracentrifuge. Clin Chem 18: 499– 502, 1972.

48. Matthews DR, Hosker JP, Rudenski AS, Naylor BA, Treacher DF, Turner RC. Homeostasis model assessment: insulin resistance and β-cell function from fasting plasma glucose and insulin concentrations in man. Diabetologia 28: 412–419, 1985. doi: 10.1007/BF00280883.

49. Van Dijk TH, Laskewitz AJ, Grefhorst A, Boer TS, Bloks VW, Kuipers F, Groen AK, Reijngoud DJ. A novel approach to monitor glucose metabolism using stable isotopically labelled glucose in longitudinal studies in mice. Lab Anim 47: 79–88, 2013. doi: 10.1177/0023677212473714.

50. Mencke R, Al Ali L, de Koning M-SLY, Pasch A, Minnion M, Feelisch M, van Veldhuisen DJ, van der Horst ICC, Gansevoort RT, Bakker SJL, de Borst MH, van Goor H, van der Harst P, Lipsic E, Hillebrands J-L. Serum Calcification Propensity Is Increased in Myocardial Infarction and Hints at a Pathophysiological Role Independent of Classical Cardiovascular Risk Factors. Arterioscler Thromb Vasc Biol 0, 2024. doi: 10.1161/ATVBAHA.124.320974.

51. McKenna HT, O’Brien KA, Fernandez BO, Minnion M, Tod A, McNally BD, West JA, Griffin JL, Grocott MP, Mythen MG, Feelisch M, Murray AJ, Martin DS. Divergent trajectories of cellular bioenergetics, intermediary metabolism and systemic redox status in survivors and non-survivors of critical illness. Redox Biol 41: 101907, 2021. doi: 10.1016/j.redox.2021.101907.

52. Bialkowska A, Ghaleb A, Nandan M, Yang V. Improved Swiss-Rolling Technique For Intestinal Tissue Preparation For Immunohistochemical And Immunofluorescent Analyses. J Vis Exp 113: e54161, 2016. doi: 10.3791/54161.

53. Mopuri R, Kalyesubula M, Rosov A, Edery N, Moallem U, Dvir H. Improved Folch Method for Liver-Fat Quantification. Front Vet Sci 7, 2021. doi: 10.3389/fvets.2020.594853.

54. Horscroft JA, Burgess SL, Hu Y, Murray AJ. Altered Oxygen Utilisation in Rat Left Ventricle and Soleus after 14 Days, but Not 2 Days, of Environmental Hypoxia. PLOS ONE 10: e0138564, 2015. doi: 10.1371/journal.pone.0138564.

55. Sowton AP, Padmanabhan N, Tunster SJ, McNally BD, Murgia A, Yusuf A, Griffin JL, Murray AJ, Watson ED. *Mtrr* hypomorphic mutation alters liver morphology, metabolism and fuel storage in mice. Mol Genet Metab Rep 23: 100580, 2020. doi: 10.1016/j.ymgmr.2020.100580.

56. Bligh EG, Dyer WJ. A Rapid Method of Total Lipid Extraction and Purification. Can J Biochem Physiol 37: 911–917, 1957. doi: 10.1139/o59-099.

57. McNally BD, Ashley DF, Hänschke L, Daou HN, Watt NT, Murfitt SA, MacCannell ADV, Whitehead A, Bowen TS, Sanders FWB, Vacca M, Witte KK, Davies GR, Bauer R, Griffin JL, Roberts LD. Long-chain ceramides are cell non-autonomous signals linking lipotoxicity to endoplasmic reticulum stress in skeletal muscle. Nat Commun 13: 1748, 2022. doi: 10.1038/s41467-022-29363-9.

58. Hall Z, Chiarugi D, Charidemou E, Leslie J, Scott E, Pellegrinet L, Allison M, Mocciaro G, Anstee QM, Evan GI, Hoare M, Vidal-Puig A, Oakley F, Vacca M, Griffin JL. Lipid Remodeling in Hepatocyte Proliferation and Hepatocellular Carcinoma. Hepatology 73: 1028–1044, 2021. doi: 10.1002/hep.31391.

59. Smith CA, Want EJ, O’Maille G, Abagyan R, Siuzdak G. XCMS: Processing Mass Spectrometry Data for Metabolite Profiling Using Nonlinear Peak Alignment, Matching, and Identification. Anal Chem 78: 779–787, 2006. doi: 10.1021/ac051437y.

60. Sud M, Fahy E, Cotter D, Brown A, Dennis EA, Glass CK, Merrill AH, Murphy RC, Raetz CRH, Russell DW, Subramaniam S. LMSD: LIPID MAPS structure database. Nucleic Acids Res 35: D527–532, 2007. doi: 10.1093/nar/gkl838.

61. Kleiner DE, Brunt EM, Van Natta M, Behling C, Contos MJ, Cummings OW, Ferrell LD, Liu Y-C, Torbenson MS, Unalp-Arida A, Yeh M, McCullough AJ, Sanyal AJ, Network NSCR. Design and validation of a histological scoring system for nonalcoholic fatty liver disease. Hepatology 41: 1313–1321, 2005. doi: 10.1002/hep.20701.

62. Taylor SR, Ramsamooj S, Liang RJ, Katti A, Pozovskiy R, Vasan N, Hwang S-K, Nahiyaan N, Francoeur NJ, Schatoff EM, Johnson JL, Shah MA, Dannenberg AJ, Sebra RP, Dow LE, Cantley LC, Rhee KY, Goncalves MD. Dietary fructose improves intestinal cell survival and nutrient absorption. Nature 597: 263–267, 2021. doi: 10.1038/s41586-021-03827-2.

63. Schindelin J, Arganda-Carreras I, Frise E, Kaynig V, Longair M, Pietzsch T, Preibisch S, Rueden C, Saalfeld S, Schmid B, Tinevez J-Y, White DJ, Hartenstein V, Eliceiri K, Tomancak P, Cardona A. Fiji: an open-source platform for biological-image analysis. Nat Methods 9: 676–682, 2012. doi: 10.1038/nmeth.2019.

64. Hippisley-Cox J, Coupland C, Brindle P. Development and validation of QRISK3 risk prediction algorithms to estimate future risk of cardiovascular disease: prospective cohort study. BMJ 357: j2099, 2017. doi: 10.1136/bmj.j2099.

65. Pratt DS, Kaplan MM. Evaluation of Abnormal Liver-Enzyme Results in Asymptomatic Patients. N Engl J Med 342: 1266–1271, 2000. doi: 10.1056/NEJM200004273421707.

66. Larsen FJ, Schiffer TA, Borniquel S, Sahlin K, Ekblom B, Lundberg JO, Weitzberg E. Dietary Inorganic Nitrate Improves Mitochondrial Efficiency in Humans. Cell Metab 13: 149–159, 2011. doi: 10.1016/j.cmet.2011.01.004.

67. Assaad H, Yao K, Tekwe CD, Feng S, Bazer FW, Zhou L, Carroll RJ, Meininger CJ, Wu G. Analysis of energy expenditure in diet-induced obese rats. Front Biosci 19: 967–985, 2014. doi: 10.2741/4261.

68. Simonson DC, DeFronzo RA. Indirect calorimetry: methodological and interpretative problems. Am J Physiol Endocrinol Metab 258: E399–412, 1990. doi: 10.1152/ajpendo.1990.258.3.E399.

69. Stump CS, Henriksen EJ, Wei Y, Sowers JR. The metabolic syndrome: Role of skeletal muscle metabolism. Ann Med 38: 389–402, 2006. doi: 10.1080/07853890600888413.

70. Jansson E. On the significance of the respiratory exchange ratio after different diets during exercise in man. Acta Physiol Scand 114: 103–110, 1982. doi: 10.1111/j.1748-1716.1982.tb06958.x.

71. Burkholder TJ, Fingado B, Baron S, Lieber RL. Relationship between muscle fiber types and sizes and muscle architectural properties in the mouse hindlimb. J Morphol 221: 177–190, 1994. doi: 10.1002/jmor.1052210207.

72. Pappachan JM, Varughese GI, Sriraman R, Arunagirinathan G. Diabetic cardiomyopathy: Pathophysiology, diagnostic evaluation and management. World J Diabetes 4: 177–189, 2013. doi: 10.4239/wjd.v4.i5.177.

73. Bugger H, Abel ED. Molecular mechanisms of diabetic cardiomyopathy. Diabetologia 57: 660–671, 2014. doi: 10.1007/s00125-014-3171-6.

74. Yin FC, Spurgeon HA, Rakusan K, Weisfeldt ML, Lakatta EG. Use of tibial length to quantify cardiac hypertrophy: application in the aging rat. Am J Physiol Heart Circ Physiol 243: H941–H947, 1982. doi: 10.1152/ajpheart.1982.243.6.H941.

75. Warren S. The effect of insulin on pathologic glycogen deposits in Diabetes mellitus. Am J Med Sci 179: 482–488, 1930. doi: 10.1097/00000441-193004000-00003.

76. Reichelt ME, Mellor KM, Curl CL, Stapleton D, Delbridge LMD. Myocardial glycophagy — A specific glycogen handling response to metabolic stress is accentuated in the female heart. J Mol Cell Cardiol 65: 67–75, 2013. doi: 10.1016/j.yjmcc.2013.09.014.

77. Lopaschuk GD, Ussher JR, Folmes CD, Jaswal JS, Stanley WC. Myocardial fatty acid metabolism in health and disease. Physiol Rev 90: 207–258, 2010. doi: 10.1152/physrev.00015.2009.

78. Neubauer S. The Failing Heart — An Engine Out of Fuel. N Engl J Med 356: 1140–1151, 2007. doi: 10.1056/NEJMra063052.

79. Younossi ZM, Koenig AB, Abdelatif D, Fazel Y, Henry L, Wymer M. Global epidemiology of nonalcoholic fatty liver disease—Meta-analytic assessment of prevalence, incidence, and outcomes. Hepatology 64: 73–84, 2016. doi: 10.1002/hep.28431.

80. He X, Wang Y, Zhang W, Li H, Luo R, Zhou Y, Liao CL M, Huang H, Lv X, Xie Z, He M. Screening differential expression of serum proteins in AFP-negative HBV-related hepatocellular carcinoma using iTRAQ –MALDI-MS/MS. Neoplasma 61: 17–26, 2014. doi: 10.4149/neo_2014_001.

81. Zou X, Zhang D, Song Y, Liu S, Long Q, Yao L, Li W, Duan Z, Wu D, Liu L. HRG switches TNFR1-mediated cell survival to apoptosis in Hepatocellular Carcinoma. Theranostics 10: 10434– 10447, 2020. doi: 10.7150/thno.47286.

82. Sanders FWB, Acharjee A, Walker C, Marney L, Roberts LD, Imamura F, Jenkins B, Case J, Ray S, Virtue S, Vidal-Puig A, Kuh D, Hardy R, Allison M, Forouhi N, Murray AJ, Wareham N, Vacca M, Koulman A, Griffin JL. Hepatic steatosis risk is partly driven by increased de novo lipogenesis following carbohydrate consumption. Genome Biol 19: 79, 2018. doi: 10.1186/s13059-018-1439-8.

83. McNally B, Griffin JL, Roberts LD. Dietary inorganic nitrate: From villain to hero in metabolic disease? Mol Nutr Food Res 60: 67–78, 2016. doi: 10.1002/mnfr.201500153.

84. Larsen FJ, Schiffer TA, Ekblom B, Mattsson MP, Checa A, Wheelock CE, Nyström T, Lundberg JO, Weitzberg E. Dietary nitrate reduces resting metabolic rate: a randomized, crossover study in humans. Am J Clin Nutr 99: 843–850, 2014. doi: 10.3945/ajcn.113.079491.

85. Chiappini F, Coilly A, Kadar H, Gual P, Tran A, Desterke C, Samuel D, Duclos-Vallée J-C, Touboul D, Bertrand-Michel J, Brunelle A, Guettier C, Le Naour F. Metabolism dysregulation induces a specific lipid signature of nonalcoholic steatohepatitis in patients. Sci Rep 7: 46658, 2017. doi: 10.1038/srep46658.

86. Hernández-Alvarez MI, Sebastián D, Vives S, Ivanova S, Bartoccioni P, Kakimoto P, Plana N, Veiga SR, Hernández V, Vasconcelos N, Peddinti G, Adrover A, Jové M, Pamplona R, Gordaliza-Alaguero I, Calvo E, Cabré N, Castro R, Kuzmanic A, Boutant M, Sala D, Hyotylainen T, Orešič M, Fort J, Errasti-Murugarren E, Rodrígues CMP, Orozco M, Joven J, Cantó C, Palacin M, Fernández-Veledo S, Vendrell J, Zorzano A. Deficient Endoplasmic Reticulum-Mitochondrial Phosphatidylserine Transfer Causes Liver Disease. Cell 177: 881–895.e17, 2019. doi: 10.1016/j.cell.2019.04.010.

87. Roberts LD, Ashmore T, McNally BD, Murfitt SA, Fernandez BO, Feelisch M, Lindsay R, Siervo M, Williams EA, Murray AJ, Griffin JL. Inorganic Nitrate Mimics Exercise-Stimulated Muscular Fiber-Type Switching and Myokine and γ-Aminobutyric Acid Release. Diabetes 66: 674– 688, 2017. doi: 10.2337/db16-0843.

88. Milsom AB, Fernandez BO, Garcia-Saura MF, Rodriguez J, Feelisch M. Contributions of Nitric Oxide Synthases, Dietary Nitrite/Nitrate, and Other Sources to the Formation of NO Signaling Products. *Antioxid Redox Signal* 17: 422–432, 2012. doi: 10.1089/ars.2011.4156.

89. Pannala AS, Mani AR, Spencer JPE, Skinner V, Bruckdorfer KR, Moore KP, Rice-Evans CA. The effect of dietary nitrate on salivary, plasma, and urinary nitrate metabolism in humans. Free Radic Biol Med 34: 576–584, 2003. doi: 10.1016/S0891-5849(02)01353-9.

90. Bryda EC. The Mighty Mouse: The Impact of Rodents on Advances in Biomedical Research. Mo Med 110: 207–211, 2013.

91. Carlström M, Persson AEG, Larsson E, Hezel M, Scheffer PG, Teerlink T, Weitzberg E, Lundberg JO. Dietary nitrate attenuates oxidative stress, prevents cardiac and renal injuries, and reduces blood pressure in salt-induced hypertension. Cardiovasc Res 89: 574–585, 2011. doi: 10.1093/cvr/cvq366.

92. Li X, Zhuge Z, Carvalho LRRA, Braga VA, Lucena RB, Li S, Schiffer TA, Han H, Weitzberg E, Lundberg JO, Carlström M. Inorganic nitrate and nitrite ameliorate kidney fibrosis by restoring lipid metabolism via dual regulation of AMP-activated protein kinase and the AKT-PGC1α pathway. Redox Biol 51: 102266, 2022. doi: 10.1016/j.redox.2022.102266.

93. Gee LC, Massimo G, Lau C, Primus C, Fernandes D, Chen J, Rathod KS, Hamers AJP, Filomena F, Nuredini G, Ibrahim AS, Khambata RS, Gupta AK, Moon JC, Kapil V, Ahluwalia A. Inorganic nitrate attenuates cardiac dysfunction: roles for xanthine oxidoreductase and nitric oxide. Br J Pharmacol 179: 4757–4777, 2022. doi: 10.1111/bph.15636.

94. Bhaswant M, Brown L, McAinch AJ, Mathai ML. Beetroot and Sodium Nitrate Ameliorate Cardiometabolic Changes in Diet-Induced Obese Hypertensive Rats. Mol Nutr Food Res 61: 1700478, 2017. doi: 10.1002/mnfr.201700478.

95. Richter K, Konzack A, Pihlajaniemi T, Heljasvaara R, Kietzmann T. Redox-fibrosis: Impact of TGFβ1 on ROS generators, mediators and functional consequences. Redox Biol 6: 344–352, 2015. doi: 10.1016/j.redox.2015.08.015.

96. Liu Z-Y, Liu Z-Y, Lin L-C, Song K, Tu B, Zhang Y, Yang J-J, Zhao J-Y, Tao H. Redox homeostasis in cardiac fibrosis: Focus on metal ion metabolism. Redox Biol 71: 103109, 2024. doi: 10.1016/j.redox.2024.103109.

97. Schopfer FJ, Cipollina C, Freeman BA. Formation and Signaling Actions of Electrophilic Lipids. Chem Rev 111: 5997–6021, 2011. doi: 10.1021/cr200131e.

98. Schopfer FJ, Khoo NKH. Nitro-Fatty Acid Logistics: Formation, Biodistribution, Signaling, and Pharmacology. Trends Endocrinol Metab 30: 505–519, 2019. doi: 10.1016/j.tem.2019.04.009.

99. Kansanen E, Bonacci G, Schopfer FJ, Kuosmanen SM, Tong KI, Leinonen H, Woodcock SR, Yamamoto M, Carlberg C, Ylä-Herttuala S, Freeman BA, Levonen A-L. Electrophilic Nitro-fatty Acids Activate NRF2 by a KEAP1 Cysteine 151-independent Mechanism. Journal of Biological Chemistry 286: 14019–14027, 2011. doi: 10.1074/jbc.M110.190710.

100. Sanyal A, Poklepovic A, Moyneur E, Barghout V. Population-based risk factors and resource utilization for HCC: US perspective. Curr Med Res Opin 26: 2183–2191, 2010. doi: 10.1185/03007995.2010.506375.

101. Anstee QM, Reeves HL, Kotsiliti E, Govaere O, Heikenwalder M. From NASH to HCC: current concepts and future challenges. Nat Rev Gastroenterol Hepatol 16: 411–428, 2019. doi: 10.1038/s41575-019-0145-7.

102. Sawa T, Ohshima H. Nitrative DNA damage in inflammation and its possible role in carcinogenesis. Nitric Oxide 14: 91–100, 2006. doi: 10.1016/j.niox.2005.06.005.

103. Masarone M, Rosato V, Dallio M, Gravina AG, Aglitti A, Loguercio C, Federico A, Persico M. Role of Oxidative Stress in Pathophysiology of Nonalcoholic Fatty Liver Disease. Oxid Med Cell Longev 2018: 9547613, 2018. doi: 10.1155/2018/9547613.

104. Khalifi S, Rahimipour A, Jeddi S, Ghanbari M, Kazerouni F, Ghasemi A. Dietary nitrate improves glucose tolerance and lipid profile in an animal model of hyperglycemia. Nitric Oxide 44: 24–30, 2015. doi: 10.1016/j.niox.2014.11.011.

105. Huang PL. eNOS, metabolic syndrome and cardiovascular disease. Trends Endocrinol Metab 20: 295–302, 2009. doi: 10.1016/j.tem.2009.03.005.

106. Tousoulis D, Papageorgiou N, Androulakis E, Siasos G, Latsios G, Tentolouris K, Stefanadis C. Diabetes Mellitus-Associated Vascular Impairment: Novel Circulating Biomarkers and Therapeutic Approaches. J Am Coll Cardiol 62: 667–676, 2013. doi: 10.1016/j.jacc.2013.03.089.

107. Matthews VB, Hollingshead R, Koch H, Croft KD, Ward NC. Long-Term Dietary Nitrate Supplementation Does Not Prevent Development of the Metabolic Syndrome in Mice Fed a High-Fat Diet. Int J Endocrinol 2018: 1–8, 2018. doi: 10.1155/2018/7969750.

108. Santamaria P. Nitrate in vegetables: toxicity, content, intake and EC regulation. J Sci Food Agric 86: 10–17, 2006. doi: 10.1002/jsfa.2351.

109. Bondonno NP, Pokharel P, Bondonno CP, Erichsen DW, Zhong L, Schullehner J, Frederiksen K, Kyrø C, Hendriksen PF, Hodgson JM, Dalgaard F, Blekkenhorst LC, Raaschou-Nielsen O, Sigsgaard T, Dahm CC, Tjønneland A, Olsen A. Source-specific nitrate intake and all-cause mortality in the Danish Diet, Cancer, and Health Study. Eur J Epidemiol, 2024. doi: 10.1007/s10654-024-01133-5

