## Supplementary material for "Chronic inorganic nitrate supplementation does not improve metabolic health and worsens disease progression in mice with diet-induced obesity": Suppl. Material

**Affiliations:**
[1] Department of Physiology, Development and Neuroscience, University of Cambridge, Cambridge, CB2 3EG, UK
[2] Department of Biochemistry and Systems Biology Centre, University of Cambridge, Cambridge, CB2 1GA, UK
[3] Clinical & Experimental Sciences, Faculty of Medicine, University of Southampton, Southampton, SO16 6YD UK
[4] Welcome Trust-MRC Institute of Metabolic Science Metabolic Research Laboratories, Addenbrooke's Hospital, Hills Road, Cambridge CB2 0QQ, UK
[5] The Rowett Institute, University of Aberdeen, Foresterhill, Aberdeen, AB25 2ZD, UK

### Supplementary Methods

#### *Study Design – Tissue Collection*

Following an overnight fast 5 (± 2) days prior to the end of the treatment period, blood was sampled from the lateral tail vein and fasted blood glucose immediately quantified (Accu-Chek Compact glucometer, Roche). The remaining blood was rapidly centrifuged (4000 x *g*, 10 min, 4°C; $K_3EDTA$ anticoagulant), plasma collected and snap frozen for clinical chemistry analysis.

Following terminal anaesthetization of mice and collection of the terminal blood sample, the heart was rapidly excised, atria and extraneous tissue removed, and the heart weighed, before the apex was removed and placed into ice-cold biopsy-preservation media (BIOPS: 2.77 mM $CaK_2EGTA$, 7.23 mM $K_2EGTA$, 6.56 mM $MgCl_2$, 50 mM MES, 5.77 mM ATP, 15 mM PCr, 20 mM imidazole, 20 mM taurine, 0.5 mM dithiothreitol, pH 7.1). A transverse mid-section 1-2 mm above the apex was carefully sectioned and placed into ice-cold paraformaldehyde (4% in phosphate-buffered saline; #J61899, Alfa-Asaer, Heysham, UK) for subsequent histological analysis, before the remaining left ventricle was separated and snap frozen. The liver was removed from the abdominal cavity and the left lateral lobe (LLL) quickly divided into 3 equal sections; one section was placed into ice-cold BIOPS, one section placed in neutral buffered formalin (#11699455, VWR International, Lutterworth, UK) and the remaining third snap frozen for molecular analyses. The right lateral lobe was fixed in isopentane pre-cooled in dry ice.

Following removal of the heart and liver, the carcass was weighed, nose-to-anus length measured, and body composition determined by dual energy x-ray absorptiometry (Lunar PIXImus II, GE Medical Systems Ltd.). Body composition parameters were calculated using PIXImus software (Lunar Corporation, Madison, WI, USA) following exclusion of the skull, per the manufacturer's instructions. Following the DEXA scan the right soleus and gastrocnemius muscles were dissected and placed into ice-cold BIOPS.

*Metabolic Cages*

Mice were acclimatized to indirect calorimetry cages (Promethion®, Sable Systems, Germany) for 24-hours prior to data collection. Cages were housed in a thermostatic, light-cycling cabinet (CAB-16, Sable Systems) to maintain a constant temperature, humidity and photoperiod equivalent to home cages throughout the experimental period.

Each mouse had *ad libitum* access to food and water of their relevant dietary intervention throughout their time in the metabolic cages, which was continuously measured via MM-2 load cells (Promethion®, Sable Systems) pre-calibrated to known masses. Indirect calorimetry was achieved through a pull-mode air flow generator (Promethion Core™ CGF, Sable Systems), calibrated to wet and dry air and zero to 5000 ppm carbon dioxide. Air was pulled through each metabolic cage at 2 $L.min^{-1}$, with subsamples analyzed for oxygen, carbon dioxide and water vapor allowing measurement of $\dot{V}O_2$ and $\dot{V}CO_2$ from each mouse.

Following the 48-hour protocol in the metabolic cages, mice underwent a body composition scan using time domain nuclear magnetic resonance (TD-NMR; LF50H Minispec, Bruker, Coventry, UK) to allow metabolic cage data to be corrected body mass covariates, as appropriate. Data were acquired using minispec software (v3.0, connected to OPUS v7.0, both Bruker).

*Histology*

Picrosirius red (PSR) staining for collagen was carried out using 0.1% Direct Red in saturated picric acid (12 $g.L^{-1}$; #A2520, AppliChem, Darmstadt, Germany) for 1h. Periodic acid Schiff (PAS) staining for glycogen was carried out using 0.5% periodic acid (5 min) and Schiff's reagent (10 min) with nuclei counter stained with haematoxylin. To correct for any PAS staining not due to glycogen deposition, one section for each sample was treated with 0.5% α-amylase (type VI-B from porcine pancreas, ~2.5 U/ml) for 20 min prior to staining.

For cryosectioning of fix-frozen liver, tissue was embedded in optimum cutting temperature compound (#KMA-0100-00A, CellPath) and sectioned at 10 μm in a cryostat (OTFAS-001, Bright Instruments, Luton, UK) at -20°C. Sections were allowed to adhere to slides at room temperature for 10 min before sections were fixed in ice-cold formalin for 10 min. Neutral lipid was then stained in oil red O (ORO; 0.3% (*w/v*) in 60% isopropanol) for 20 min. Nuclei were counterstained with haematoxylin. Following staining, the slides were mounted in glycerin gelatin at 60°C and cover slips sealed with clear nail varnish once the mountant had set.

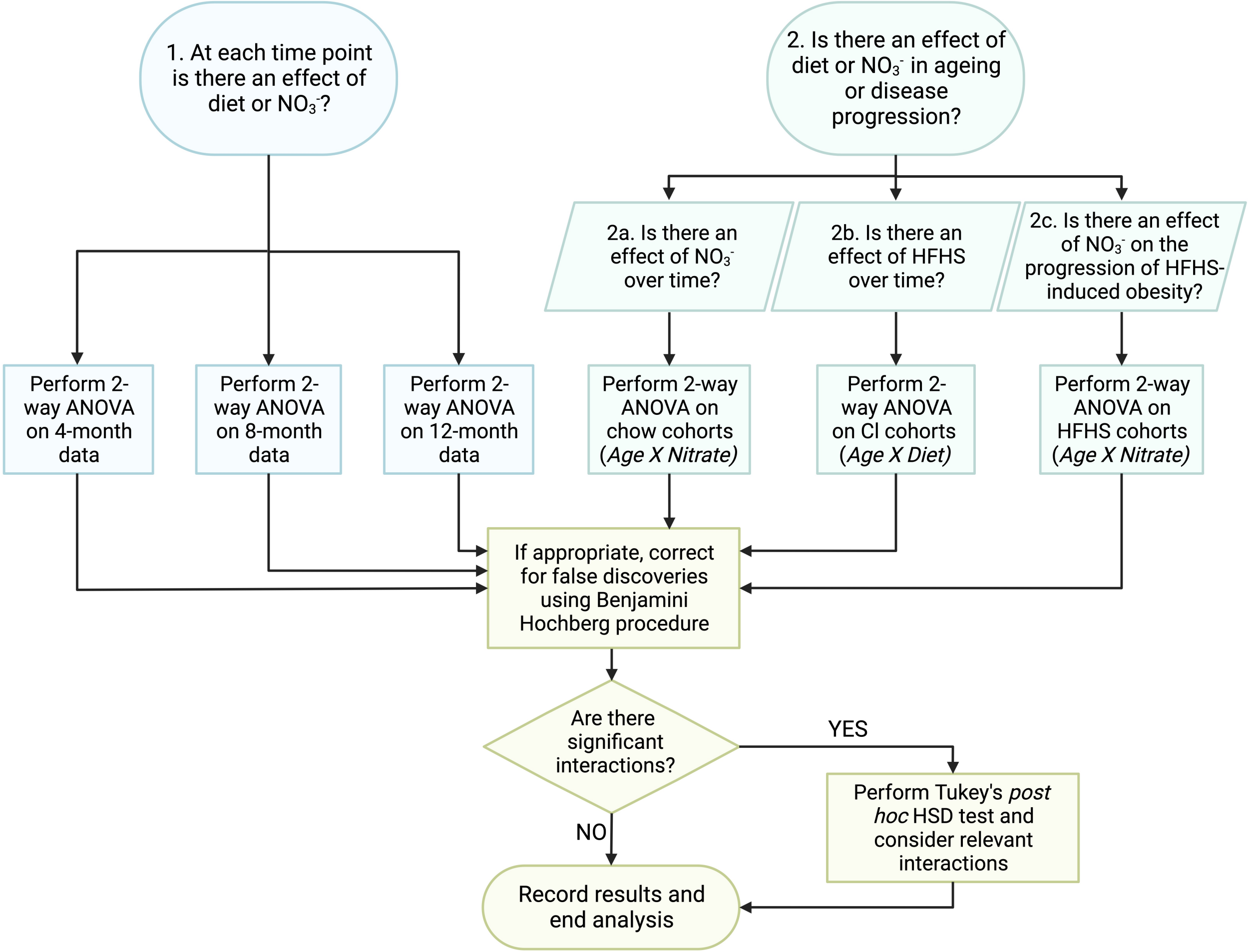

1. At each time point is there an effect of diet or NO3-?
2. Is there an effect of diet or NO3- in ageing or disease progression?
2a. Is there an effect of NO3- over time?
2b. Is there an effect of HFHS over time?
2c. Is there an effect of NO3- on the progression of HFHS-induced obesity?
Perform 2-way ANOVA on 4-month data
Perform 2-way ANOVA on 8-month data
Perform 2-way ANOVA on 12-month data
Perform 2-way ANOVA on chow cohorts (Age X Nitrate)
Perform 2-way ANOVA on CI cohorts (Age X Diet)
Perform 2-way ANOVA on HFHS cohorts (Age X Nitrate)
If appropriate, correct for false discoveries using Benjamini Hochberg procedure
Are there significant interactions?
YES
NO
Perform Tukey's post hoc HSD test and consider relevant interactions
Record results and end analysis

**Supplementary Figure 1.** Statistical workflow used for data analysis. Created with BioRender.com.

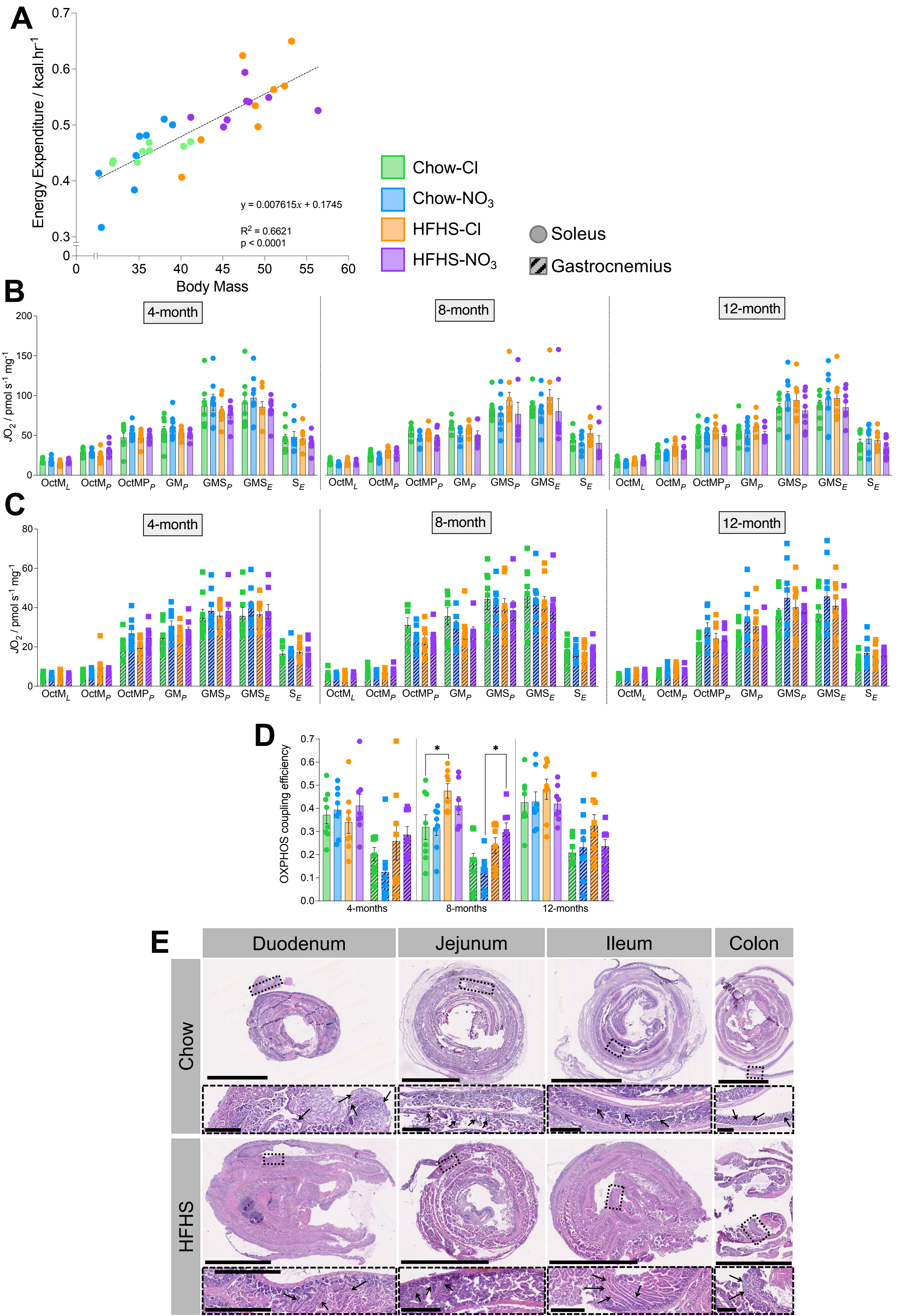

A
Energy Expenditure / kcal.hr$^{-1}$
Body Mass
y = 0.007615x + 0.1745
$R^2$ = 0.6621
p < 0.0001
Chow-Cl
Chow-$NO_3$
HFHS-Cl
HFHS-$NO_3$
Soleus
Gastrocnemius
B
4-month
8-month
12-month
$JO_2$ / pmol s$^{-1}$ mg$^{-1}$
$OctM_L$ $OctM_P$ $OctMP_P$ $GM_P$ $GMS_P$ $GMS_E$ $S_E$
C
4-month
8-month
12-month
$JO_2$ / pmol s$^{-1}$ mg$^{-1}$
$OctM_L$ $OctM_P$ $OctMP_P$ $GM_P$ $GMS_P$ $GMS_E$ $S_E$
D
OXPHOS coupling efficiency
4-months
8-months
12-months
E
Duodenum
Jejunum
Ileum
Colon
Chow
HFHS

**Supplementary Figure 2.** Energy expenditure is unaffected by HFHS diet, but intestinal villi are longer in HFHS-fed mice.

(A) Energy expenditure (calculated *via* the Weir equation) as a function of total body mass in 10-month-old mice. Line was fitted with non-linear regression to compare if the experimental groups fit to the same line. N=7-8 / group.

(B) Mitochondrial respiratory function assessed via high-resolution respirometry in saponin-permeabilised soleus muscle fibres. Data represent mean ± SEM; N=7-8 / group.

(C) Mitochondrial respiratory function assessed via high-resolution respirometry in saponin-permeabilised gastrocnemius muscle fibres. Data represent mean ± SEM; N=7-8 / group.

(D) The proportion of OXPHOS not limited by LEAK respiration, supported by octanoyl-carnitine and malate in saponin-permeabilised skeletal muscle fibres (OXPHOS coupling efficiency). Data represent mean ± SEM; N=7-8 / group. $^{*}p<0.05$; two-way ANOVA with Tukey's *post hoc* HSD test for multiple comparisons.

(E) Representative histological images of intestinal segments stained with H&E from mice fed chow or HFHS diets for 8-weeks. Scale bars on overview images represent 2.5 mm, and on 10x magnified inserts (dashed boxes) represent 250 µm. Examples of intact villi/crypts used for quantification are shown with arrows on the inserts.

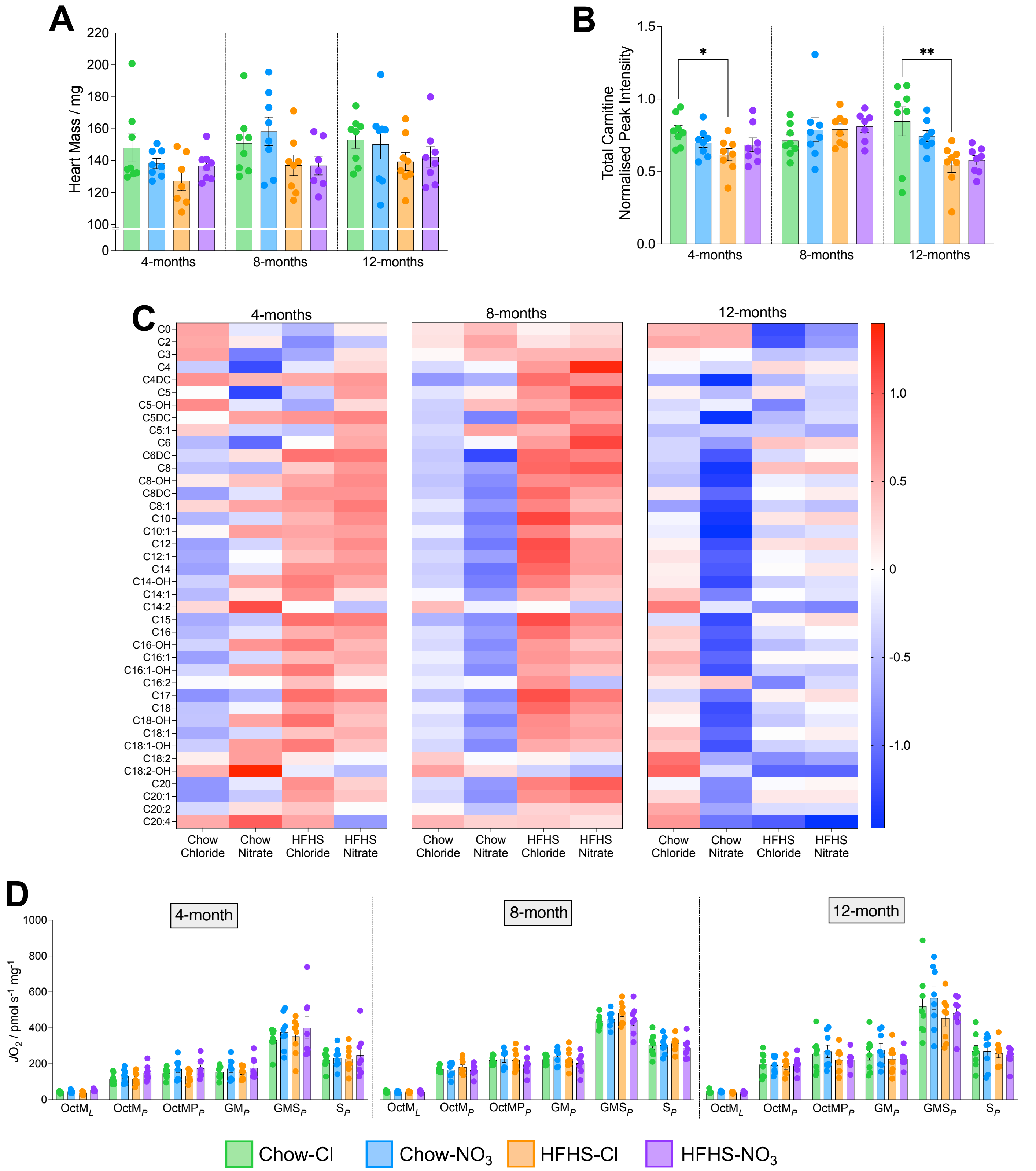

A
Heart Mass / mg
4-months
8-months
12-months
B
Total Carnitine Normalised Peak Intensity
*
**
C
4-months
8-months
12-months
C0
C2
C3
C4
C4DC
C5
C5-OH
C5DC
C5:1
C6
C6DC
C8
C8-OH
C8DC
C8:1
C10
C10:1
C12
C12:1
C14
C14-OH
C14:1
C14:2
C15
C16
C16-OH
C16:1
C16:1-OH
C16:2
C17
C18
C18-OH
C18:1
C18:1-OH
C18:2
C18:2-OH
C20
C20:1
C20:2
C20:4
Chow Chloride
Chow Nitrate
HFHS Chloride
HFHS Nitrate
D
4-month
8-month
12-month
$JO_2$ / pmol s$^{-1}$ mg$^{-1}$
$OctM_L$
$OctM_P$
$OctMP_P$
$GM_P$
$GMS_P$
$S_P$
Chow-Cl
Chow-$NO_3$
HFHS-Cl
HFHS-$NO_3$

**Supplementary Figure 3.** Alterations to acyl-carnitines and mitochondrial function in hearts from HFHS-fed and $NO_3$-suplemented mice.

(A) Gross heart mass across all experimental groups.

(B) Total cardiac acyl-carnitine content, determined by targeted liquid chromatography-mass spectrometry, measured as total peak area ratio, normalised to an appropriate internal standards and sample protein concentration. $^{*}p<0.05$, $^{**}p<0.01$; two-way ANOVA with Tukey's *post hoc* HSD test for multiple comparisons.

(C) Relative concentration of individual free and acyl-carnitines across experimental groups. Cell colour represents mean concentration for each group. Data determined from targeted liquid chromatography-mass spectrometry analysis, with peak area ratios normalised to an appropriate internal standard and sample protein concentration. Data was autoscaled for presentation on the same heatmap.

(D) Mitochondrial respiratory function assessed via high-resolution respirometry in saponin-permeabilised cardiac muscle fibres.

Data represent mean ± SEM; N=7-8 / group.

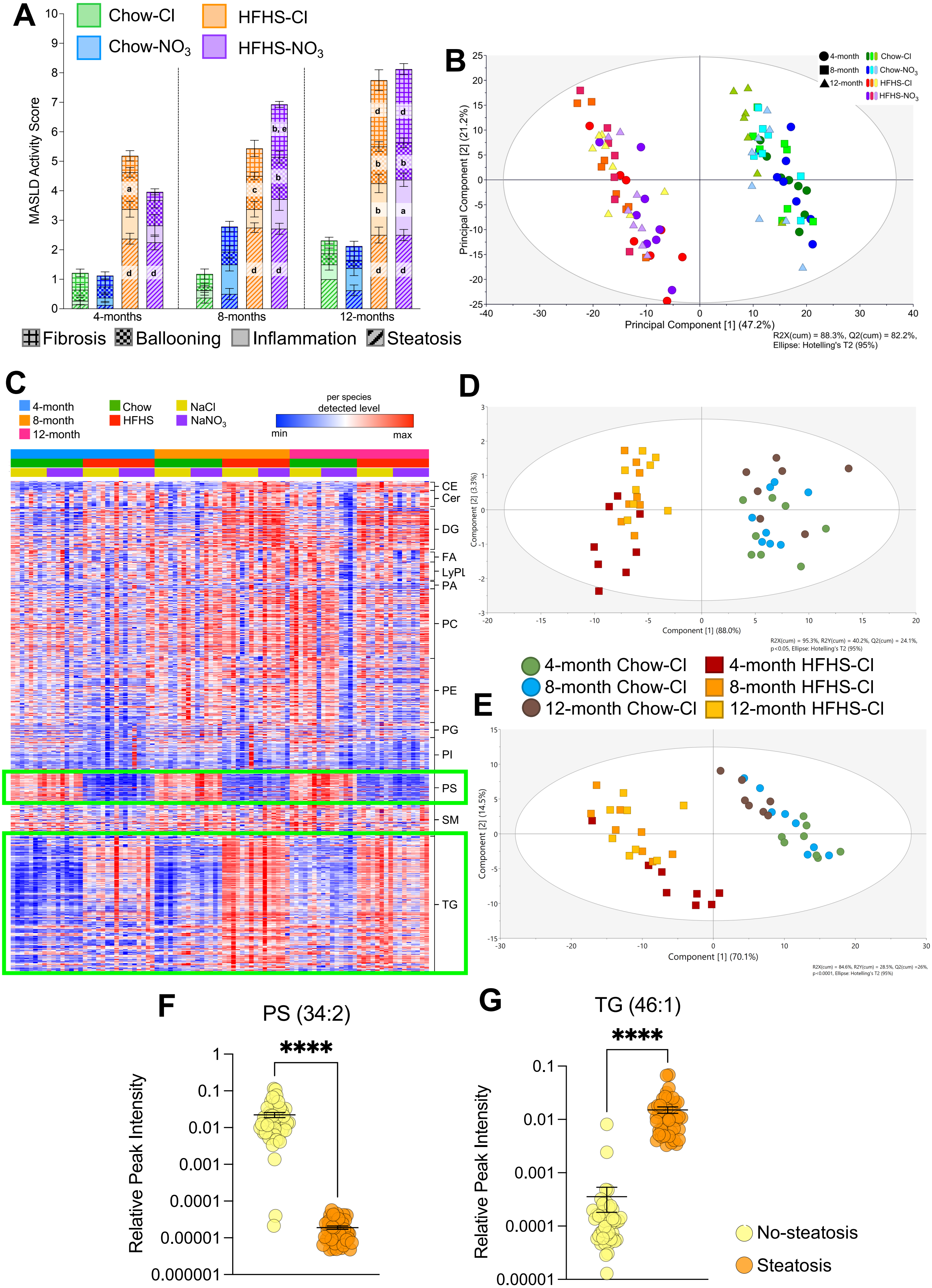
A
Chow-Cl
HFHS-Cl
Chow-NO3
HFHS-NO3
MASLD Activity Score
4-months
8-months
12-months
Fibrosis
Ballooning
Inflammation
Steatosis
B
4-month
8-month
12-month
Principal Component [1] (47.2%)
Principal Component [2] (21.2%)
R2X(cum) = 88.3%, Q2(cum) = 82.2%,
Ellipse: Hotelling's T2 (95%)
C
4-month
8-month
12-month
Chow
HFHS
NaCl
NaNO3
per species
detected level
min
max
CE
Cer
DG
FA
LyPL
PA
PC
PE
PG
PI
PS
SM
TG
D
Component [1] (88.0%)
Component [2] (3.3%)
4-month Chow-Cl
8-month Chow-Cl
12-month Chow-Cl
4-month HFHS-Cl
8-month HFHS-Cl
12-month HFHS-Cl
E
Component [1] (70.1%)
Component [2] (14.5%)
F
PS (34:2)
Relative Peak Intensity
G
TG (46:1)
Relative Peak Intensity
No-steatosis
Steatosis

**Supplementary Figure 4.** HFHS-feeding is associated with hepatic lipidome remodelling.

(A) Individual components of MASLD activity score. $^{a}p<0.05$, $^{b}p<0.01$, $^{c}p<0.001$, $^{d}p<0.0001$, compared with nitrate-matched chow-fed mice at the same time point; $^{e}p<0.05$, compared with HFHS-Cl mice at the same time point; two-way ANOVA with Tukey's *post hoc* HSD test for multiple comparisons. Data represent mean ± SEM.

(B) Principal component analysis of the hepatic lipidome detected by untargeted liquid chromatography-mass spectrometry. Data was log transformed and Pareto scaled prior to analysis; $R^2X_{(cum)} = 88.3\%$, $Q^2_{(cum)} = 82.2\%$.

(C) Heat-map of all lipids detected *via* open-profiling lipidomics grouped by experimental condition and lipid species. Plot generated using Morpheus software (Broad Institute). Differences in phosphatidylserines (PS) and triacylglycerols (TG) highlighted.

(D) Partial least squares-discriminant analysis (PLS-DA) of the PS complement of the hepatic lipidome of chloride-supplemented mice only. Data was log transformed and Pareto scaled prior to analysis; $R^2X_{(cum)} = 95.3\%$, $R^2Y_{(cum)} = 40.2\%$, $Q^2_{(cum)} = 24.1\%$, $p<0.05$.

(E) PLS-DA of the TG complement of the hepatic lipidome of chloride-supplemented mice only. Data was log transformed and Pareto scaled prior to analysis; $R^2X_{(cum)} = 84.6\%$, $R^2Y_{(cum)} = 28.5\%$, $Q^2_{(cum)} = 26.0\%$, $p<0.0001$.

For (A)-(E), all data represent N=7-8 / group.

(F) Relative levels of PS(34:2) in livers from mice with no histologically defined steatosis (MAS steatosis component scored 0 or 1) and those with histologically defined steatosis (MAS steatosis component scored 2 or 3).

(G) Relative levels of TG(46:1) in livers from mice with no histologically defined steatosis (MAS steatosis component scored 0 or 1) and those with histologically defined steatosis (MAS steatosis component scored 2 or 3).

For (F)-(G), data represent mean ± SEM; N=44-49 / group; $^{****}p<0.0001$.

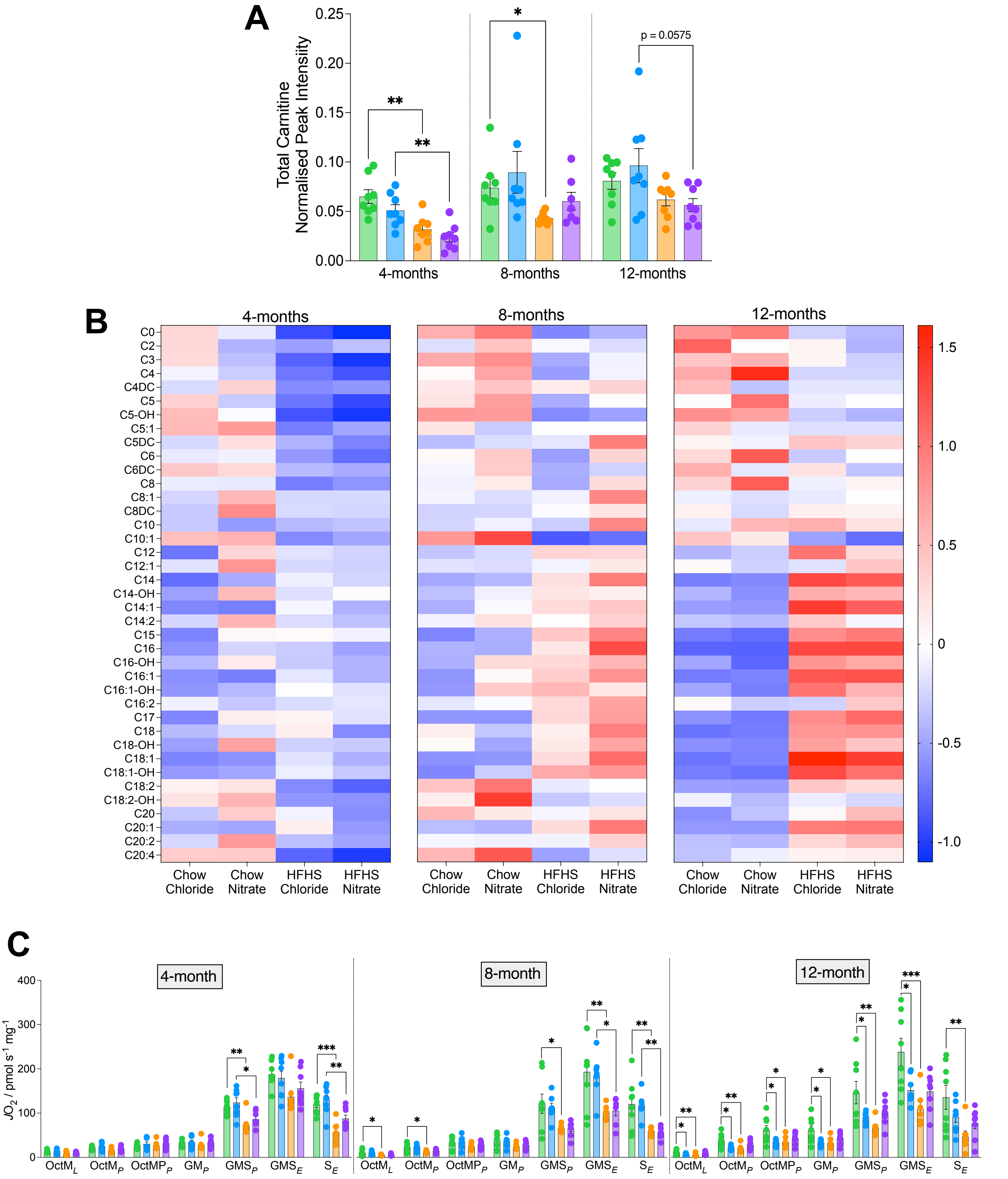

A
Total Carnitine Normalised Peak Intensity
0.25
0.20
0.15
0.10
0.05
0.00
4-months
8-months
12-months
**
*
p = 0.0575
B
4-months
8-months
12-months
C0
C2
C3
C4
C4DC
C5
C5-OH
C5:1
C5DC
C6
C6DC
C8
C8:1
C8DC
C10
C10:1
C12
C12:1
C14
C14-OH
C14:1
C14:2
C15
C16
C16-OH
C16:1
C16:1-OH
C16:2
C17
C18
C18-OH
C18:1
C18:1-OH
C18:2
C18:2-OH
C20
C20:1
C20:2
C20:4
Chow Chloride
Chow Nitrate
HFHS Chloride
HFHS Nitrate
1.5
1.0
0.5
0
-0.5
-1.0
C
4-month
8-month
12-month
JO2 / pmol s-1 mg-1
400
300
200
100
0
OctML
OctMP
OctMPP
GMP
GMSP
GMSE
SE
***

**Supplementary Figure 5.** Alterations to acyl-carnitines and mitochondrial function in livers from HFHS-fed and $NO_3$-suplemented mice.

(A) Total hepatic acyl-carnitine content, determined by targeted liquid chromatography-mass spectrometry, measured as total peak area ratio, normalised to an appropriate internal standards and sample protein concentration. $^{*}p<0.05$, $^{**}p<0.01$; two-way ANOVA with Tukey's post hoc HSD test for multiple comparisons.

(B) Relative concentration of individual free and acyl-carnitines across experimental groups. Cell colour represents mean concentration for each group. Data determined from targeted liquid chromatography-mass spectrometry analysis, with peak area ratios normalised to an appropriate internal standard and sample protein concentration. Data was autoscaled for presentation on the same heatmap.

(C) Mitochondrial respiratory function assessed *via* high-resolution respirometry in liver homogenate. $^{*}p<0.05$, $^{**}p<0.01$, $^{***}p<0.001$; two-way ANOVA with Tukey's *post hoc* HSD test for multiple comparisons.

Data represent mean ± SEM; N=7-8 / group.

### SUPPLEMENTARY TABLES

**Supplementary Table 1.** Detail of primers used for RT-qPCR.

| **Gene** | **Tissue**: Liver (L) or Heart (H) | **Catalogue Number** (GeneGlobe ID) | **Transcript Reference** (NCBI) |
|---|---|---|---|
| *Acadl* | L | QT00101248 | NM_007381 |
| *Acc1* | L | QT01554441 | NM_133360 |
| *Actb* | H | QT00095242 | NM_007393 |
| *Akr1b10* | L | QT00130361 | NM_172398 |
| *Fasn* | L | QT00149240 | NM_007988 |
| *Gsk3a* | H | QT01202866 | NM_001031667 |
| *Gys1* | H | QT00162099 | NM_030678 |
| *Hadh* | L | QT00147672 | NM_008212 |
| *Hmbs* | L | QT00494130 | NM_013551 |
| *Hrg* | L | QT00172249 | NM_053176 |
| *Pdgfb* | L | QT00266910 | NM_011057 |
| *Ppp1ca* | H | QT00104055 | NM_031868 |
| *Rn18s* | H, L | QT02448075 | NR_003278 |
| *Scd1* | L | QT00291151 | NM_009127 |
| *Serpine1* | L | QT00154756 | NM_008871 |
| *Spp1* | L | QT00157724 | NM_009263 |
| *Srsf4* | L | QT01056132 | NM_020587 |
| *Ywhaz* | H | QT00105350 | NM_011740 |

Details of QuantiTect primer IDs and transcript references in the NCBI sequence used for gene expression analysis. All primers were purchased from QIAGEN.

**Table 2.** Semi-quantitative histological scoring system for determining MASLD activity score (MAS).

| **Feature** | **Stain** | **Magnification** | **Description** | **Score** |
|---|---|---|---|---|
| Steatosis | ORO | x20 | <5% | 0 |
| | | | 5 – 33% | 1 |
| | | | >33 – 66% | 2 |
| | | | >66% | 3 |
| Hepatocyte ballooning | H&E | x10 | None | 0 |
| | | | Few balloon cells | 1 |
| | | | Many cells/prominent ballooning | 2 |
| Lobular inflammation | H&E | x20 | No foci | 0 |
| | | | <2 foci per field | 1 |
| | | | 2 – 4 foci per field | 2 |
| | | | >4 foci per field | 3 |
| Fibrosis | PSR | x10 | None | 0 |
| | | | Mild perisinusoidal | 1a* |
| | | | Moderate perisinusoidal | 1b* |
| | | | Portal/periportal | 1c* |
| | | | Perisinusoidal and poral/periportal | 2 |
| | | | Bridging fibrosis | 3 |
| | | | Cirrhosis | 4 |

ORO = Oil red O; H&E = haematoxylin and eosin; PSR = picrosirius red. *For statistical analysis sections graded 1a were given a score of 0.5, 1b a score of 1 and 1c a score of 1.5.
